# Nuclear Argonaute:miRNA complexes recognize target sequences within chromatin and silence gene expression

**DOI:** 10.1101/2025.05.08.652285

**Authors:** Cristina Hofman, Jiaxin Hu, Rut Bryl, Victor Tse, David R. Corey

## Abstract

RNA interference (RNAi) in mammalian cells involves recognition of mRNA in the cytoplasm and inhibition of translation. Both protein RNAi factors and miRNAs, however, are present in mammalian cell nuclei. It is unclear how this nuclear localization affects endogenous gene expression. Here, we use chimeric eCLIP to identify complexes of Argonaute 2 (AGO2) and miRNAs. We identify the most abundant miRNAs associated with chromatin and their chromatin-associated RNA targets. Chimeric eCLIP revealed that *High mobility group AT-Hook A* (*HMGA2*) was the most compelling target for miRNA-mediated gene binding. There are four confirmed *let-7* miRNA sites within the 3’-UTR in the cytoplasm or nucleus and three within chromatin-associated RNA. The expression of mature HMGA2 mRNA was repressed by let-7 in the cytoplasm and nucleus. *let-7* had little effect splicing or transcription. Our data validate chimeric eCLIP as a powerful method for experimentally identifying promising miRNA:RNA interactions. Rather than a solely cytoplasmic event, binding of RNAi factors to mRNA targets may begin in the nucleus through a mechanism that can reduce RNA levels in both the cytoplasm and the nucleus. miRNA-mediated silencing of mRNAs may be influenced by both nuclear and cytoplasmic interactions.

## Introduction

RNA interference (RNAi) is a versatile mechanism of gene regulation (1–3). At the heart of RNAi, an Argonaute (AGO) protein is loaded with a microRNA (miRNA) or synthetic duplex RNA to form the RNA induced silencing complex (RISC). In the cytoplasm, RISC acts as a programmable regulatory factor, with the small RNA facilitating recognition of an RNA target by Watson-Crick base pairing. The AGO protein protects the small RNA, promotes binding to target messenger RNA (mRNA) and silences these targets through either enzymatic cleavage or recruitment of additional proteins for translational repression and transcript degradation (4–7). The diversity of synthetic RNAs is essentially infinite and there are hundreds of well validated miRNAs in human cells, suggesting the potential to target almost any accessible cellular RNA.

In mammalian cells, RNAi is generally associated with the silencing of genes in the cytoplasm. However, miRNAs (8) and protein RNAi factors like AGO proteins (9–12) are also present and functional in mammalian cell nuclei. Exogenous synthetic RNAs have been shown to modulate transcription (13–18) and splicing (19–21). Endogenous miRNAs have been found to control transcription (14, 22–24), splicing (25, 26), silence mobile transposons in quiescent cells (27), and influence gene expression in spermatogenic cells through association with meiotic chromatin (28). In other organisms, including *S. pombe* and *A. thaliana*, endogenous RISC has roles in regulation of chromatin dynamics (29–31) in addition to its role in transcriptional regulation. While these studies suggest the potential for diverse and robust nuclear regulation in mammalian cells, the full biological impact of nuclear RNAi remains to be understood.

While miRNAs have hundreds of potential targets inside human cells (32) and may produce many statistically significant effects on gene expression, target identification is rarely a simple task (33, 34). Our previous work found that AGO binding in a gene 3’UTR does not always lead to repression (35) and other studies have shown that single miRNAs are not essential for viability or development (36, 37). These findings suggest that commonly used methods of target identification, including genetic manipulation of miRNAs or CLIP to identify AGO binding sites, may produce false positive results and fail to identify promising candidates. While bioinformatic target predictions can be used to identify seed sequence complementarity in an unbiased manner, careful analysis must be done to keep up with evolving stringencies in the definition of functional miRNAs (33, 38–40). Ultimately, the identity and impact of regulatory miRNAs likely varies among cell types and finding the biologically significant miRNA:target interaction among the many potential interactions can be like finding a needle in a haystack (41).

Fortunately, contemporary experimental tools allow for higher confidence identification of some miRNA target interactions. While previous work largely relied on genetic manipulation and microarrays (38,39,44), advances in crosslinking and immunoprecipitation (CLIP) allow us to identify AGO2 binding sites in an unbiased fashion (43–45). More specialized CLIP techniques even allow for the identification of specific miRNA:mRNA interactions (48,49). While beneficial advances, these techniques still have their limitations and offer no guarantee that they will identify all biologically relevant control points for miRNAs. However, in tandem, they offer powerful tools for prioritizing potential miRNA targets for experimental validation.

In this report we use chimeric eCLIP (48) to obtain complementary datasets that define potential miRNA targets in HCT116 human colorectal cancer cells. These data identify *high mobility group AT-Hook A* (*HMGA2*) as the best candidate for miRNA-mediated regulation. We demonstrate that recognition of the *HMGA2* 3’-untranslated region begins when the RNA is associated with chromatin and can also be detected throughout cell nuclei and the cytoplasm. Gene silencing can be observed in the nucleus. For *HMGA2*, however, changes in transcription and splicing are not observed. These data suggests that the journey of miRNA-mediated recognition and silencing can begin in mammalian cell nuclei and chromatin.

## Materials and Methods

### Cell culture

Wild-type HCT116, DROSHA^-/-^, NLS-AGO2, and HT-29 cells were cultured in McCoy’s 5A medium supplemented with 10% FBS. HeLa cells were grown in Dulbecco’s Modified Eagle’s medium (DMEM) supplemented with 10% FBS. All cells were grown at 37°C in 5% CO_2_ and passed when 70-80% confluent.

### Preparation of cytoplasm, nucleus, nucleoplasm, and chromatin extracts

Cells were seeded at 2.5 x 10^6^ cells per dish into 150 mm dishes and supplemented with 60 mL of fresh media on day 2 after seeding. Cells were harvested 3 days after seeding with trypsin. Subcellular fractionation to isolate cytoplasm and nucleus extracts were as previously described and isolation of cytoplasm, nucleoplasm and chromatin extracts was as previously described with modifications (11, 49, 50). Samples were kept on ice and ice-cold buffers were used. Briefly, harvested cells were pelleted and washed with PBS. Cells were pelleted and lysed with hypotonic lysis buffer (HLB) (10 mM Tris HCl pH 7.4, 10 mM NaCl, 3 mM MgCl_2_, 2.5% NP-40, 0.5 mM DTT). Nuclei were pelleted and the supernatant was collected as the cytoplasmic fraction. Nuclei were washed a total of six times with HLB. After washing, the nuclei were lysed with modified Wuarin-Schibler buffer (MWS) (10 mM Tris HCl pH 7.0, 4 mM EDTA, 0.3 M NaCl, 1 M urea, 1% NP-40). Chromatin was pelleted and supernatant was collected as the nucleoplasm fraction. Chromatin was washed twice with MWS and lysed by sonication in lysis buffer (50 mM Tris HCl pH 7.4, 120 mM NaCl, 0.05% NP-40, 1 mM EDTA, 1 mM DTT).

### Western blot analysis

Proteins were separated on 4–20 % gradient Mini-PROTEAN® TGXTM precast gels (Bio-Rad). After gel electrophoresis, proteins were wet transferred to nitrocellulose membrane (0.45 μM, GE Healthcare Life Sciences) at 100 V for 90 min. Membranes were blocked for 1 hr. at room temperature with 5% milk in 1× PBS containing 0.05% Tween-20. Blocked membranes were incubated with the primary antibodies in blocking buffer at 4°C on rocking platform overnight: anti-AGO1 1:1000 (Cell Signaling, #5053), anti-AGO2 1:2000 (Fujifilm Wako, 015-22031), anti-AGO3 1:1000 (Cell Signaling, #5054), anti-AGO4 1:1000 (Cell Signaling, #6913), anti-TNRC6A 1:5000 (A302-329A, Bethyl), anti-DROSHA 1:500 (Cell Signaling, #3364), anti-Dicer (Cell Signaling, #3363), anti-Calnexin 1:5000 (Cell Signaling, #2433), anti-β-tubulin 1:5000 (Sigma-Aldrich, T5201), SP1 1:5000 (Cell Signaling, #5931) Histone H3 1:20K (Cell Signaling, #2650). After primary antibody incubation, membranes were washed 3 × 10 min at room temperature with 1 × PBS + 0.05% Tween-20 (PBST 0.05%) and then incubated for 1 h at room temperature with respective secondary antibodies in blocking buffer. Membranes were washed again 3 × 15 min in PBST 0.05%. Washed membranes were soaked with HRP substrate according to manufacturer’s recommendations (SuperSignal™ West Pico Chemiluminescent substrate, Thermo Scientific) and exposed to films. The films were scanned, and bands were quantified using ImageJ software.

### Co-immunoprecipitation

Cell lysates were precleared for 1 hour at 4°C in IP dilution buffer (20 mM Tris HCl pH 7.4, 150mM NaCl, 3mM MgCl_2_), then incubated overnight with specific antibodies. Complexes were recovered with 80µL of protein G Dynabeads (Invitrogen), then washed five times with IP wash buffer (20 mM Tris HCl pH 7.4, 400mM NaCl, 3mM MgCl_2_, 0.05% NP-40, 0.01% SDS) (1mL, 5 min at 4°C). Protein was eluted with 20µL of 2X Tris-SDS loading buffer incubating for 15 minutes at RT and transferred to a new tube for western blot.

### miR-eCLIP

Cells were seeded in 150 mm dishes and grown until 80-90% confluent. Cells were then UV-crosslinked on ice at 254nm (400mJ/cm^2^). Cells were collected or fractionated as described above. Prior to storage, cell pellets and lysates were flash frozen.

For miR-eCLIP experiments, the standard eCLIP protocol (45) was modified to enable chimeric ligation of miRNA and mRNA (48). Studies were performed by Eclipse Bioinnovations Inc. (SanDiego, www.eclipsebio.com). Cells were then lysed in 1000 µL of eCLIP lysis mix and sonicated (QSonica Q800R2) for 5 minutes, 30 seconds on / 30 seconds off with an energy setting of 75% amplitude. The chromatin fractions from UV crosslinked cells were lysed in 1000 µL of eCLIP lysis mix and sonicated (QSonica Q800R2) for 17 minutes, 30 seconds on / 30 seconds off with an energy setting of 75% amplitude. The cytoplasmic fractions from UV crosslinked cells were sonicated (QSonica Q800R2) for 5 minutes, 30 seconds on / 30 seconds off with an energy setting of 75% amplitude and diluted in 2X volume of 1.5X eCLIP lysis buffer. Equivalent amounts of sample lysates were digested with RNase-I (Ambion). A primary mouse monoclonal AGO2/EIF2C2 antibody (sc-53521, Santa Cruz Biotechnology) was incubated for 1 hour with magnetic beads pre-coupled to the secondary antibody (M-280 Sheep Anti-Mouse IgG Dynabeads, Thermo Fisher 11202D) and added to the homogenized lysate for overnight immunoprecipitation at 4°C. Following overnight IP, 2% of the sample was taken as the paired size-matched input with the remainder magnetically separated and washed with eCLIP high stringency wash buffers. The chromatin samples were subjected to DNase treatment (TURBO DNase Thermo Fisher) followed by eCLIP high stringency wash step. Chimeric ligation was then performed on-bead at room temperature for 1 hour with T4 RNA ligase (NEB). IP samples were then dephosphorylated with alkaline phosphatase (FastAP, Thermo Fisher) and T4 PNK (NEB) and an RNA adapter was ligated to the 3’ ends. IP and input samples were cut from the membrane at the AGO2 protein band size to 75kDa above. Western blot was visualized using anti-AGO2 primary antibody (50683-RP02, SinoBiological) at a 1:4000 dilution, with TrueBlot anti-rabbit secondary antibody (18-8816-31, Rockland) at 1:8000 dilution. RNA adapter ligation, reverse transcription, DNA adapter ligation, and PCR amplification were performed as previously described.

After sequencing, samples were processed with Eclipsebio’s proprietary analysis pipeline (v1). UMIs were pruned from read sequences using umi_tools (v1.1.1). Next, 3’ adapters were trimmed from reads using cutadapt (v3.2). Reads were then mapped to a custom database of repetitive elements and rRNA sequences. All non-repeat mapped reads were mapped to the hg38 genome using STAR (v2.7.7a). PCR duplicates were removed using umi_tools (v1.1.1). AGO2 eCLIP peaks were identified within eCLIP samples using the peak caller CLIPper (v2.0.1). For each peak, IP versus input fold enrichments and p-values were calculated.

miRNAs from miRBase (v22.1) were “reverse mapped” to any reads that did not map to repetitive elements or the genome using bowtie (v1.2.3). The miRNA portion of each read was then trimmed, and the remainder of the read was mapped to the genome using STAR (v2.7.7a). PCR duplicates were resolved using umi_tools (v1.1.1), and miRNA target clusters were identified using CLIPper (v2.0.1). Each cluster was annotated with the names of miRNAs responsible for that target. Peaks were annotated using transcript information from GENCODE v41 with the following priority hierarchy to define the final annotation of overlapping features: protein coding transcript (CDS, UTRs, intron), followed by non-coding transcripts (exon, intron).

### Transfection of miRNA inhibitors and mimics

miRNA mimics were ordered from IDT. The miRCURY LNA^TM^ miRNA inhibitors were purchased from QIAGEN. All transfections used Lipofectamine RNAiMAX (Invitrogen). Cells were seeded into six-well plates 24 hours before transfection, at 300K cells per well for wild type HCT116 cells and 450K for DROSHA-/- cells. At the next day, 50 nM of miRNA mimics or inhibitors were transfected into cells as previously described (50). 24 hours later, cells were replaced with full culture media. Cells were harvest for qPCR 48 hours after transfection.

### RT-qPCR

Total RNA was extracted from cells using TRIzol reagent. Reverse transcription was performed using high-capacity reverse transcription kit (Applied Biosystems) per the manufacturer’s protocol. 2.0 μg of total RNA was used per 20 μL of reaction mixture. qPCR was performed on performed on a CFX96 Touch real-time PCR system (Bio-Rad) using iTaq SYBR Green Supermix (Bio-Rad). PCR reactions were done in triplicates at 55°C 2 min, 95°C 3 min and 95°C 30 s, 60°C 30 s for 40 cycles in an optical 96-well plate. The primers were listed in Supplementary Table S1. Data were normalized relative to the level of an internal control gene *RPL19*.

### RNAPII ChIP-qPCR

Cells were crosslinked with 1% formaldehyde at RT for 10 min and crosslinking was quenched with glycine. Cells were then harvested by scraping in PBS. Cells were pelleted and nuclei were isolated with hypotonic lysis buffer (HLB) (10 mM Tris HCl pH 7.4, 10 mM NaCl, 3 mM MgCl_2_, 2.5% NP-40, 0.5 mM DTT) and lysed in lysis buffer (50 mM Tris HCl pH 7.4, 120 mM NaCl, 0.05% NP-40, 1 mM EDTA, 1 mM DTT). Samples were sonicated for 20 minutes (30s on, 30s off, 50% amplitude, Qsonica Q800R3). Nuclear lysate was precleared, then incubated overnight with RNAPII antibody (05-623, Millipore Sigma, 3 µg) or normal mouse IgG (12-371, Millipore Sigma) in immunoprecipitation buffer. After antibody-protein-DNA complex was recovered with 80 µl of protein G Dynabeads (Invitrogen) for 2 hours, the beads were washed with low salt, high salt, LiCl, and TE buffers (1 mL each, 5 min at 4°C). The complex was eluted twice with 250 µl of elution buffer (1% SDS, 0.1 M NaHCO_3_) at RT. Cross-linking was reversed by adding NaCl to a final concentration of 200 mM and heating at 65°C for 2 h. Protein was digested by incubating with proteinase K at 42°C for 50 min, followed by phenol-chloroform extraction and ethanol precipitation. Samples were dissolved in nuclease free water and analyzed by qPCR. Primer sets used are listed in supplementary table 1.

### Quantitative polymerase chain reaction (qPCR) splicing assay

A qPCR splicing assay for the *HMGA2* gene was developed by adapting methods as described (51). A qPCR splicing assay was also developed for the TBP gene as an internal qPCR control. Exon-exon junction spanning primers were designed to have optimal target specificity, melting temperatures, and amplicon size. These primer sets were validated to amplify regions of interest in silico. Melt curve analysis for each primer set also demonstrates robust amplification. Each primer set used in the *HMGA2* and *TBP* qPCR splicing assay is shown in Supplemental Table 1. All primers were synthesized by Integrated DNA Technologies.

20 ng of cDNA corresponding to a given experimental condition was used as a qPCR template for the iTaq™ Universal SYBR® Green Supermix reaction (Bio-Rad Laboratories) following the manufacturer’s instructions. All reactions were set-up in 384-well qPCR plates and were run on a CFX384 Touch Real-Time PCR Detection System, followed by preliminary data analysis done in the CFX Maestro Software. As described in Harvey et al. (2021), each exon inclusion:skipping ratio was calculated using the formula: 2−ΔCt, where ΔCt = (Ct corresponding to the inclusion mRNA isoform – Ct corresponding to the skipping mRNA isoform) from the same sample.

Statistical significance in differences between experimental conditions was determined using 2-way grouped analysis of variance (ANOVA), and Tukey’s post-hoc multiple comparisons test. All statistical tests were performed in GraphPad Prism 10. Values were determined to be statistically significant if the calculated P-value was below an alpha value of ≤0.05.

## RESULTS

### Isolation of chromatin-bound RNAi factors

Previous work has detected RNAi factors in mammalian cell nuclei (9–12, 14, 16, 27, 50) and demonstrated control gene splicing and transcription (13–15, 17–26). While these data imply that regulation could occur in association with chromatin, direct observation of chromatin bound AGO or miRNA species has been complicated by competing demands during purification. Isolation of chromatin must be stringent enough to exclude cytoplasm and nucleoplasm but not so stringent as to wash away chromatin-associated AGO protein, AGO:target RNA complexes, and AGO:miRNA complexes. To address these challenges, we chose to optimize conditions for the isolation of cytoplasm and nuclear fractions in HCT116 colorectal cancer-derived cells.

We focus on HCT116 cells because we possess CRISPR/Cas9-knockout cell lines lacking critical RNAi factors and a CRISPR/Cas9-knockin AGO2-nuclear localization signal line (AGO2-NLS). Starting from published subcellular fractionation protocols (11, 49, 50), we systematically optimized pH, detergent percentage, and salt concentration of our fractionation buffers to identify conditions that prioritized separation of nucleoplasm and chromatin, while maintaining maximum protein content in the chromatin lysate. Following subcellular fractionation, we observed AGO2 protein in the chromatin fraction and confirmed the purity of the fractions using protein markers for the cytoplasm (β-tubulin), nucleoplasm (SP1), endoplasmic reticulum (calnexin), and chromatin (Histone H3) by western analysis (**Figure 1AB**).

**Figure 1.**
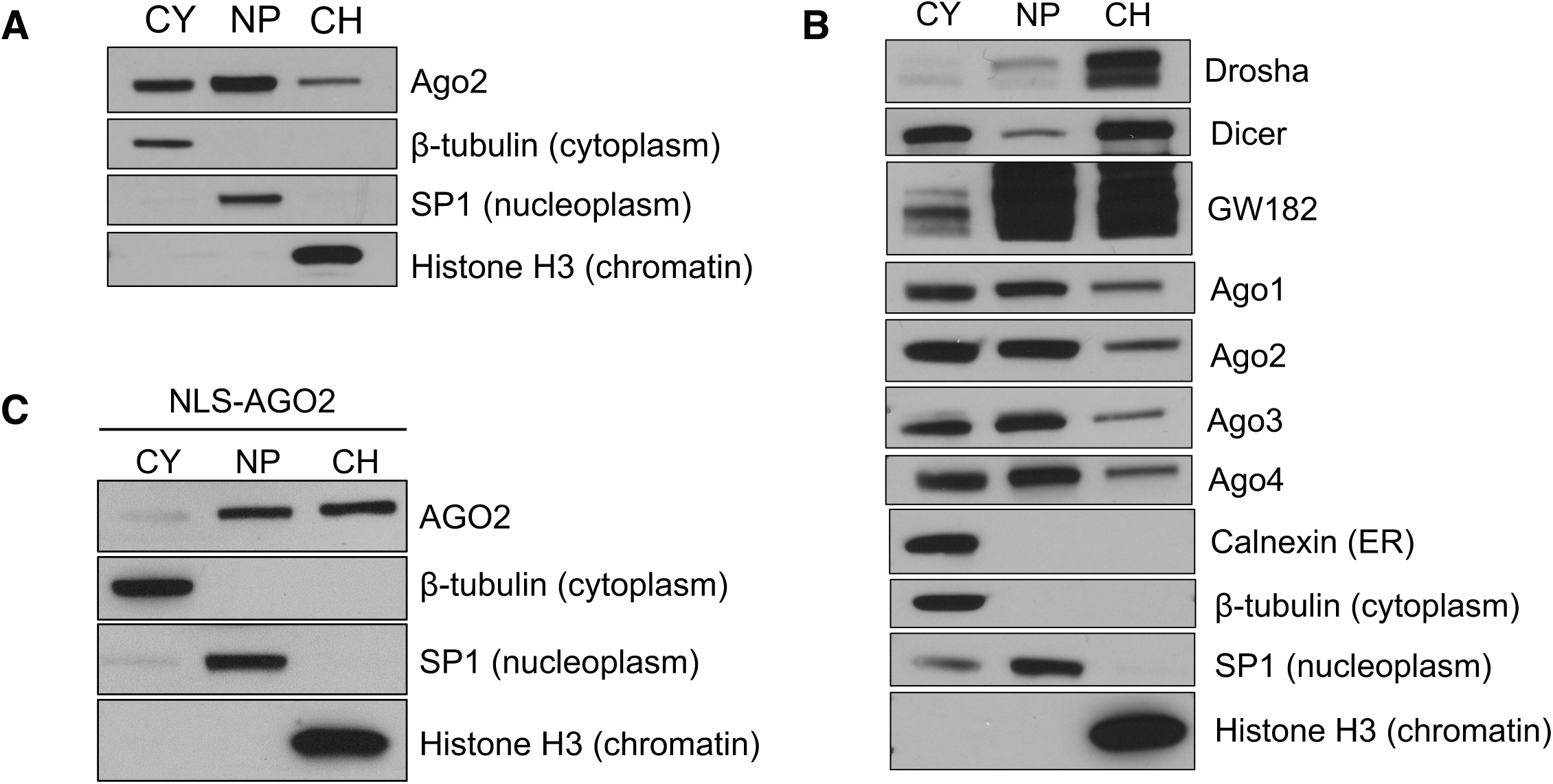
AGO2 and additional RNAi factors can associate with chromatin. (A) Western blot showing AGO2 localization and purity of cytoplasm (CY), nucleoplasm (NP) and chromatin (CH). (B) Western blot of RNAi factors and purity markers from cytoplasm, nucleoplasm, and chromatin. (C) Western blot showing AGO2 localization in NLS-AGO2 knock-in cell line

AGO has 4 paralogs in mammalian cells (AGO1-4) with AGO2 being the most abundant. AGO2 (4–6), and to a lesser extent AGO3 (52), possess the ability to catalyze the cleavage of RNA substrates. AGO1-4 bind miRNAs similarly (53). We found that AGO1-4 are present in all compartments of the cells and importantly are associated with chromatin (**Figure 1B**). Drosha and Dicer are the RNase III enzymes which catalyze the production of mature miRNAs and function in the nucleus and cytoplasm respectively. We found that Drosha is mainly present in chromatin, while Dicer is present in the cytoplasm and chromatin (**Figure 1B**), consistent with previous results (11, 54, 55).

Human GW182 or trinucleotide repeat containing protein 6A (TNRC6A) is a scaffolding protein that interacts with AGO2 and facilitates miRNA-mediated degradation of transcripts (56–62). Our data show that human TNRC6A is present in all compartments of the cell, including chromatin (**Figure 1B**), suggesting that it can act as a scaffold to promote complexes with other proteins. We note that our gel loading involves equal protein, not equal cell equivalents. This analysis is designed to evaluate purity and does not represent the absolute levels of TNRC6A or other proteins in the different chromatin, nucleoplasm, or cytoplasm relative to one another. We have not attempted to optimize recovery of cytoplasmic proteins since our focus is on chromatin and our results do not imply that there is more TNRC6A in cell nuclei relative to cytoplasm. We and others have reported previously nuclear TNRC6A (11, 16, 35, 59, 60) and used mass spectrometry to characterize its interacting protein partners in nuclear lysate (16, 56).

We used our NLS-AGO2 HCT116 cells to provide us with an alternative method for examining the nuclear interactions of AGO2 by chimeric eCLIP that is independent of subcellular fractionation. This NLS-AGO2 cell line has a nuclear localization sequence added to the N-terminus of the endogenous locus of the AGO2 gene. This construct avoids overexpression, but forces endogenous AGO2 to localize to the nucleus (50) and chromatin (**Figure 1C**). Therefore, immunoprecipitation using an anti-AGO2 antibody will isolate nuclear AGO2:RNA interactions without the need for a prior subcellular fractionation step, providing a complementary strategy for examining nuclear associations.

### Scheme for Anti-AGO2 CLEAR-CLIP and eCLIP

Interactions between miRNAs and cellular RNA targets can be predicted by computation. However, it can be difficult to sort through the many different predicted targets. To increase the likelihood that time-consuming experimental validation is focused on the most promising targets, it is useful to complement computational analysis with physical methods that detect sites where physical interactions between miRNAs, AGO proteins, and RNA targets occur.

We chose to use AGO2 chimeric enhanced crosslinking and immunoprecipitation (chimeric eCLIP) to identify these physical interactions between RISC and its RNA targets (48) (**Figure 2A**). eCLIP employs UV crosslinking prior to harvesting cells. Because the cells are still alive when the crosslinking occurs, the crosslinking reflects interactions that occur within living cells. Cellular fractions are then obtained and an AGO2 antibody is used to pull down the protein:RNA complex. To increase the likelihood of detecting direct miRNA-RNA target interactions, the immunoprecipitation is followed by a ligation step that connects a miRNA to its mRNA target.

**Figure 2.**
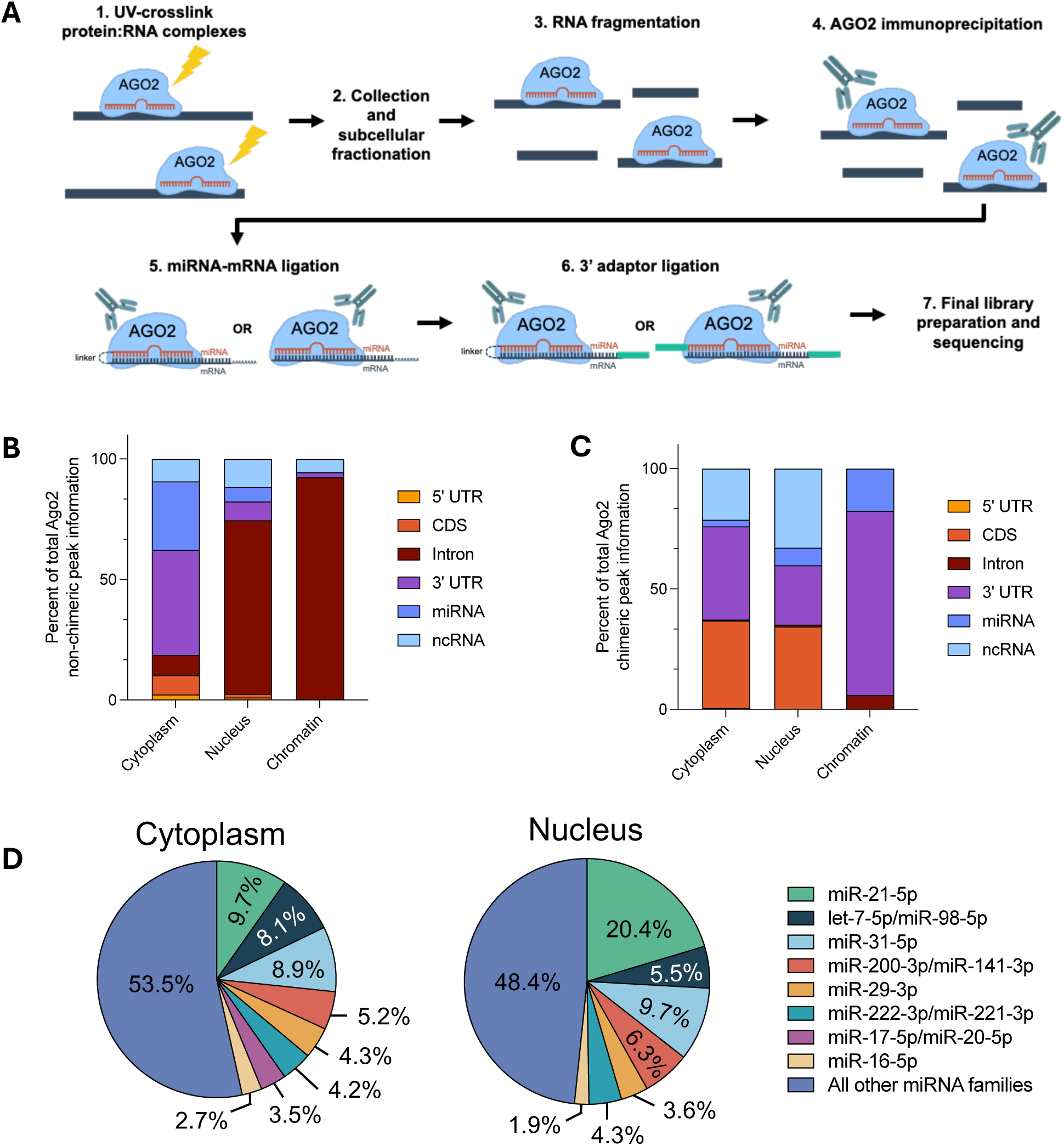
AGO2 interacts with various target RNAs in all cellular compartments. (A) Schematic outlining the chimeric eCLIP protocol used for the identification of AGO2 binding sites. Distribution of AGO2 (B) non-chimeric and (C) chimeric binding peaks across genomic features in cytoplasm, whole nucleus, and chromatin fractions. Chimeric binding peaks only include chimeric reads that include miRNAs that map to the miRGeneDB database. (D) Percentage of chimeric miRNAs from the top eight miRNA families sharing the same seed sequence.

The ligation step that joins miRNA and RNA target is not efficient and yields two groups of RNA products following digestion of the AGO2 protein. miRNA-mRNA pairs where the ligation was successful will be chimeric, with a linker connecting them. These chimeric reads identify unbiased, direct miRNA binding to targets. The remaining RNA that was bound by AGO2 will remain non-chimeric (**Figure 2A**). The non-chimeric reads, a much larger portion of the total reads, identify RNAs associate with AGO2. The method produces relatively few chimeric reads, identifying high confidence miRNA:RNA target interactions and a relatively high number of non-chimeric reads, identifying sites of AGO2 association with RNA.

To gain insight into whether functional RISC targets could be identified in chromatin, we performed two independent anti-AGO2 chimeric eCLIP experiments using whole cell lysate from wild-type HCT116 cells, whole cell lysate from AGO2-NLS HCT116 cells, wild type cytoplasm, and wild type chromatin (**Supplementary Figure 1**).

### Chimeric eCLIP: RNA target regions and miRNAs

Both total (∼56-62 million) and final non-chimeric (9.5 to 16.6 million) reads were similar in every sample regardless of whether the original was whole cell, cytoplasm, nucleus, or chromatin (**Supplementary Figure 1**). In all samples, the number of chimeric reads was much lower than the number of nonchimeric reads because the generation of chimeric reads involves a ligation step that limits the efficiency of detection. Lower detection of chimeric reads is consistent with prior results using other methods that report ligation efficiencies ranging from 1.5-5% (46). Chimeric reads were at least 10-fold lower in the chromatin sample (∼14,000 reads) relative to whole cell (∼100,000 reads), cytoplasm (∼170,000 reads), and nuclear (75,000 reads) samples.

For cytoplasmic RNA, most non-chimeric reads were associated with recognition within the 3’-untranslated region (3’-UTR) (**Figure 2B**). By contrast in the nucleus and chromatin fraction, most peaks are observed within intronic RNA, likely because of the much greater fraction of intronic RNA present prior to splicing and nuclear export. Chimeric AGO2 peaks in the cytoplasm were mainly found within the 3’-UTR (**Figure 2V**), like the non-chimeric binding peaks. By contrast to the similar results in cytoplasm for nonchimeric and chimeric AGO2 peaks, data from nuclei and chromatin showed a shift to binding within the 3’-UTR (**Figure 2C**).

Why would chimeric reads in the chromatin or nuclear fractions cluster within 3’-UTRs whereas non-chimeric reads cluster within introns? We hypothesize that the greater percentage of chimeric reads within the 3’-UTR in nucleus or chromatin (**Figure 2C**) is due to miRNA:AGO2:3’-UTR complexes being stronger and hence more detectable by the relatively less efficient chimeric eCLIP.

The chimeric reads also contain information about the specific miRNAs mediating target RNA recognition (**Figure 2D**). Using the average reads for the top 20 miRNAs in the chimeric reads from whole cell HCT116 eCLIP samples, we found that eight miRNA families make up ∼50 % of the total reads. In the cytoplasm and nucleus, these eight miRNA families account for 46.5% and 51.6% of the chimeric miRNA reads (**Figure 2D**). The number of chimeric peaks in the chromatin fraction was too small to draw conclusions about the distribution of miRNAs.

### Prioritizing genes for experimental analysis

To facilitate experimental validation, we prioritized genes that: 1) had significant numbers of chimeric reads in either cytoplasm, nucleus, or chromatin and, 2) had altered expression in wild-type versus DROSHA-/- cells. DROSHA is the most upstream enzyme in the miRNA biogenesis pathway. Knockout of the protein leads to depletion of most mature miRNAs and gene expression changes would suggest a potential miRNA-mediated effect on gene expression. From this prioritization, we chose five genes for experimental analysis, *HMGA2*, *MYC*, *ZFP36*, *ZP36L1*, and *TNFRSF12A*. All these genes have at least one chimeric peak in their 3’UTR in the cytoplasm and nucleus and *HMGA2, MYC* and *TNFRSF12A* have chimeric peaks detected in chromatin. Each gene also has non-chimeric AGO2 peaks within their 3’-UTRs of varying strengths (**Supplemental Figure 2**).

Of these gene candidates, High Mobility Group AT-Hook 2 (*HMGA2)* was the most compelling. Both non-chimeric and chimeric AGO2 peaks were detected in the 3’UTR of *HMGA2* in the cytoplasm, nucleus and chromatin (**Figure 3, Supplementary Figure 1, Supplementary Table 2**). RNAseq indicated that *HMGA2* gene expression increased in DROSHA-/- relative to WT by 5-fold, the largest positive increase of any candidate gene, suggesting a potential de-repression of miRNA-mediated silencing and consistent with canonical gene silencing mechanisms.

**Figure 3.**
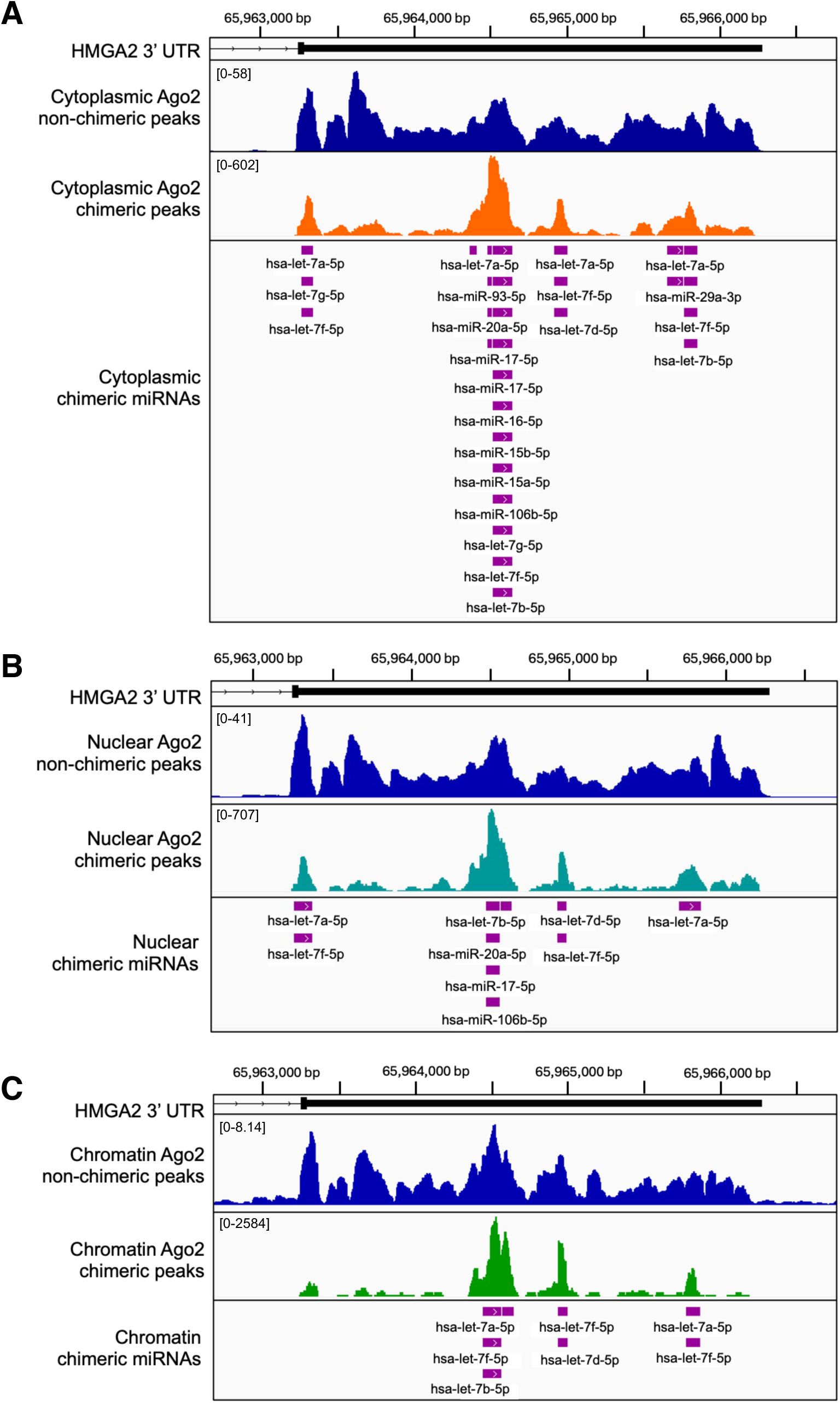
HMGA2 is bound by AGO2 in all compartments of the cell. Representative IGV browser images of chimeric eCLIP reads within the 3’UTR of HMGA2 in the (A) cytoplasm, (B) nucleus, and (C) chromatin. HMGA2 is located on chromosome 12 (chr12) and thin bars represent intronic segments of the gene while thick bars represent the exons and 3’UTR. Peak height is defined as read density in reads per million (RPM). Individual chimeric miRNA sites are separate defined AGO2 peaks with overlapping chimeric reads with a log2FC>3 (IP vs. input).

### HMGA2, a model for miRNA-mediated regulation

miRNA-mediated gene silencing is most effective when multiple miRNAs bind at adjacent sites (63–65). Binding of multiple miRNAs in proximity to one another permits cooperative interactions through the scaffolding protein TNRC6 which can bind up to three AGO proteins at one time. This allows TNRC6 to bridge three RISC complexes and stabilize their binding to target mRNA (61).

For example, within the 3’UTR of *HMGA2*, chimeric eCLIP detected >20 miRNA binding peaks in the cytoplasm and 8 miRNA binding peaks in the nucleus and chromatin (**Figure 3**). Of these binding peaks, we observed four binding sites for the *let-7* family of miRNAs using nuclear or cytoplasmic anti-AGO2 chimeric eCLIP (**Figure 4AB**), and three binding sites by chimeric eCLIP of chromatin (**Figure 4C**). Comparative analysis suggests that the *let-7* seed sequence at these binding sites are well-conserved (**Supplemental Figure 3**).

**Figure 4.**
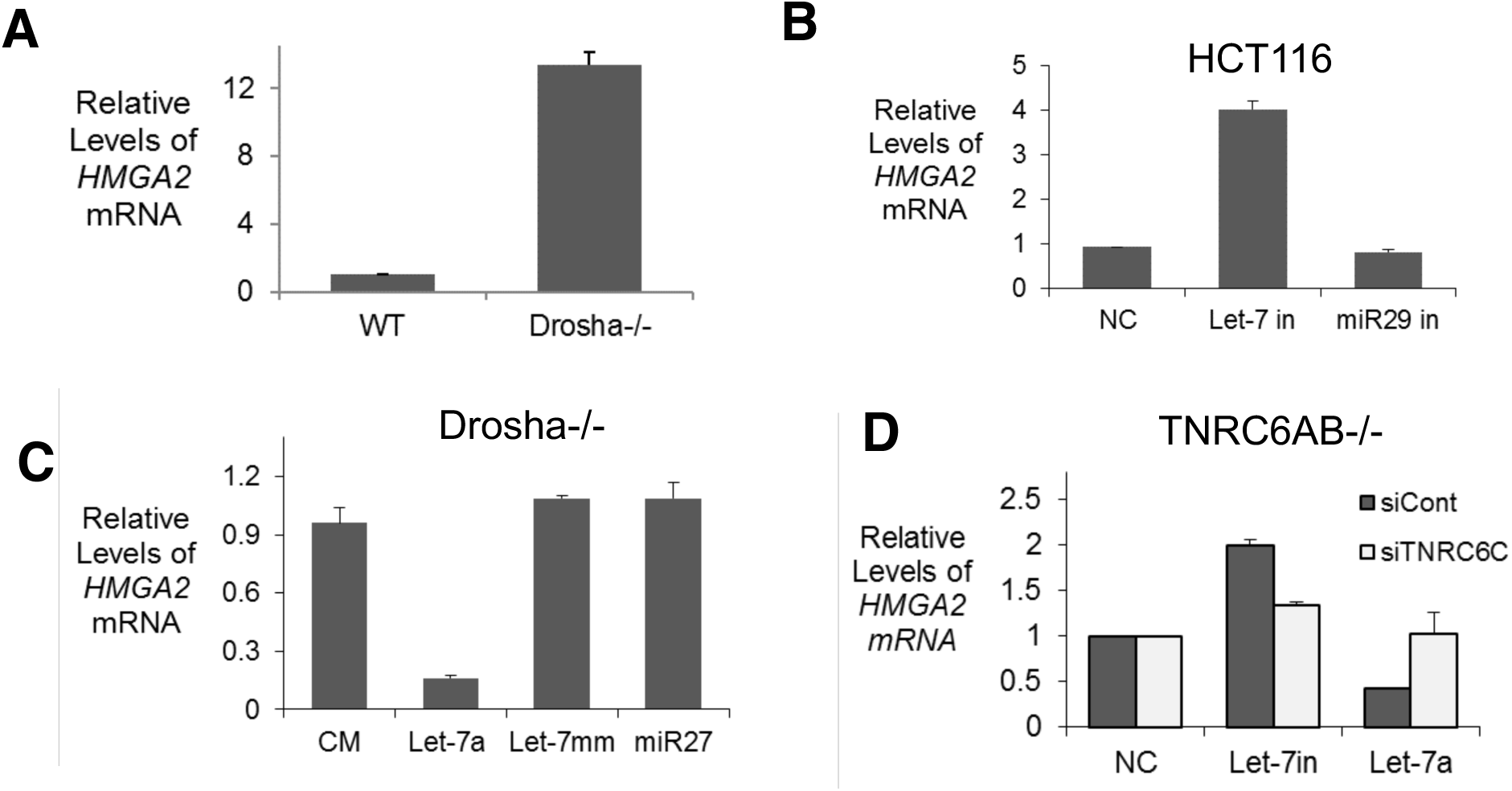
HMGA2 expression is regulated by the let-7 family of miRNAs. Relative level of mature HMGA2 mRNA in **A**) untreated WT and Drosha-/- cells, **B**) WT cells transfected with non-targeting, *let-7* family, or *miR-29* family miRNA inhibitors, and **C**) Drosha-/- cells transfected with *let-7a* or control miRNA mimics. **D**) HMGA2 mRNA level measured after knockout and knockdown of all three TNRC6 paralogs. In TNRC6A/B-/- cells, a non-targeting control and siTNRC6C siRNA were transfected, then followed by a second round of transfection of let-7 inhibitor or let-7a miRNA mimics. All values are plotted as the average of biological replicates +/-SD.

The involvement of *let-7* is consistent with previous computational and experimental analyses implicating *let-7* in the regulation of *HMGA2* expression in other cancer-derived cell lines (66–69). The *let-7* family is one of the more prevalent miRNAs detected by chimeric eCLIP (**Fig. 2D**). The combination of compelling chimeric eCLIP data and data implicating *let-7* in the control of HMGA2 in other cell types led us to focus on exploring the *let-7/HMGA2* axis as a model for understanding the contributions of nuclear and cytoplasmic interactions to gene silencing.

To find experimental support for the role of *let-7* family miRNAs in the regulation of *HMGA2* in HCT116 cells, we used complementary methods for probing miRNA-mediated regulation using RNA isolated from HCT116 whole cell lysate. As mentioned, DROSHA is the protein responsible for production of most mature miRNAs and its knockout leads to the depletion of most mature miRNAs in HCT116 cells (53). Q-PCR measurent of *HMGA2* levels showed that, relative to WT HCT116, the *HMGA2* transcript is upregulated, greater than 10-fold in *DROSHA*-/- cells (**Figure 4A**).

While analysis of the *DROSHA*-/- cells confirmed that miRNAs are involved in *HMGA2* regulation, we next wanted to determine whether the *let-7* family of miRNAs that we had identified through eCLIP play a direct role in this regulation. Synthetic oligonucleotides that are complementary to miRNAs (miRNA inhibitors or antimiRs) can block the action of miRNAs and reverse their control of gene expression (70–72). To test the effect of *let-7* miRNAs on *HMGA2* expression, we used a *let-7* miRNA family inhibitor that selectively blocks activity of the *let-7* family members in wild-type HCT116 cells. Transfection of the *let-7* family miRNA inhibitor increased *HMGA2* expression (**Figure 4B**), while a miRNA inhibitor targeting the *miR-29* family or a noncomplementary cocktail of control oligonucleotides had no effect on *HMGA2* expression (**Figure 4B**).

Duplex RNAs that mimic the sequence of *let-7* would be expected to also mimic the ability of *let-7* to repress gene expression (73). To test this hypothesis, we used DROSHA-/- HCT116 cells that lack endogenous expression of *let-7* miRNAs (53). Following transfection of a *let-7a* mimic, *HMGA2* expression was decreased relative to a noncomplementary control duplex, a *miR-27* mimic, or a mismatched duplex based on the *let-7a* sequence (**Figure 4C**). Together, these results further support that the *let-7* family of miRNAs play a direct role in the regulation of *HMGA2* expression.

TNRC6 is a scaffolding protein that interacts with AGO2 and is critical for miRNA-mediated inhibition of translation (74–79). AGO2 also retains catalytic activity that does not require TNRC6 when the guide RNA and target RNA are perfectly complementary (4–6, 80). To separate these two potential pathways of miRNA-mediated silencing, we chose to use our TNRC6AB-/- cells to examine the necessity of the TNRC6 paralogs for HMGA2 silencing. TNRC6 has 3 paralogs, and in the TNRC6AB-/- cell line, TNRC6C is upregulated (80). As a result, testing the effects of reducing expression of *TNRC6A*, *TNRCB* and *TNRC6C* is best accomplished by using an siRNA designed to reduce TNRC6C in TNRC6AB-/- cells.

When we transfected miRNA inhibitor targeting the let-7 family into *TNRC6AB-/-* cells (elevated TNRC6C expression) we observed increased *HMGA2* expression, consistent with residual *let-7* mediated gene silencing (**Figure 4D**). When TNRC6C was also silenced, *HMGA2* expression returned to control level, consistent with the TNRC6 paralogs acting together to contribute to miRNA-mediated regulation of *HMGA2*. When a *let-7a* mimic was transfected into *TNRC6AB-/-* cells, we observed a decrease in HMGA2 expression, consistent enhanced silencing by the synthetic miRNA. When *TNRC6C* was also silenced, addition of the *let-7a* mimic had no significant effect on gene expression. Our data are consistent with previous work suggesting that the TNRC6 paralogs can compensate for one another and loss of all three is necessary to observe their full impact (26, 80). Ultimately, these results suggest that HMGA2 is silenced, at least partially, through a miRNA-mediated translation repression mechanism.

### No correlation between miRNA binding and the regulation of other candidate genes

We also examined the potential for miRNA-mediated regulation of our other candidate genes, *MYC*, *ZFP36*, *ZFP36L1*, and *TNFRSF12A*. For *MYC*, we observed nonchimeric AGO2 association and chimeric *let-7* reads within the *MYC* 3’-UTR (**Supplementary Figure 4A**). The *let-7* seed sequence binding sites were relatively well conserved **(Supplementary Figure 5).** *MYC* expression was decreased in *DROSHA*-/- cells – a result opposite to what would be expected by canonical miRNA-mediated regulation (**Supplementary Figure 4B, Supplementary Table 1**).

Despite evidence of association of *let-7* within the *MYC* 3’-UTR, we observed no direct *let-7* mediated impact on gene expression. In *DROSHA-/-* cells, the transfection of *let-7a* mimics had no significant impact on the expression of *MYC* mRNA (**Supplementary Figure 3C**). In wild type cells, addition of *let-7* family miRNA inhibitors had no significant effect on *MYC* expression in cell cytoplasm or nuclei (**Supplementary Figure 3D and 3E).** We observed a similar lack of correlation between AGO2 chimeric and non-chimeric binding and regulation of gene expression at the paralog *ZFP36* and *ZFP36L1* loci (**Supplemental Figures 6 and 7**). Taken together, these data indicate that the presence of chimeric reads within a gene is no guarantee of observable biological regulation by candidate miRNAs.

### let-7 regulates *HMGA2* RNA levels in cell nuclei

After establishing an experimental foundation to support *let-7* mediated regulation of *HMGA2* expression in whole cell lysate (**Figures 3 and 4**), we investigated whether the observed strong recognition of the *HMGA2* 3’-UTR by *let-7* within the nucleus and chromatin (**Figure 3BC**) contributes to the regulation of *HMGA2* silencing. To test this hypothesis, we evaluated nuclear and cytoplasmic expression of *HMGA2* in *DROSHA*-/- cells. We also tested the impact of miRNA inhibitors and miRNA mimics on HMGA2 mRNA expression in the cytoplasm and nucleus. For comparison, we fractionated untreated WT and *DROSHA-/-* HCT116 cells and evaluated *HMGA2* expression in both the nuclear and cytoplasmic fractions.

First, we examined the impact of miRNAs in the cytoplasm, the canonical site of miRNA-mediated silencing. We observed that *HMGA2* expression was increased in the cytoplasm of *DROSHA*-/- cells relative to wild-type (**Figure 5A**). Inhibition of the *let-7* family of miRNAs led to an increase in HMGA2 expression (**Figure 5C**), and addition of a *let-7a* miRNA mimic led reduced expression of *HMGA2* in *DROSHA*-/- cells (**Figure 5E**). We also found that like in whole cell, the silencing of HMGA2 in the cytoplasm was dependent on the presence of the TNRC6 paralogs (**Figure 5G**). These results suggest that in the cytoplasm, HMGA2 is silenced through miRNA-mediated recognition of its 3’UTR by *let-7* miRNA loaded RISC and subsequent translational repression.

**Figure 5.**
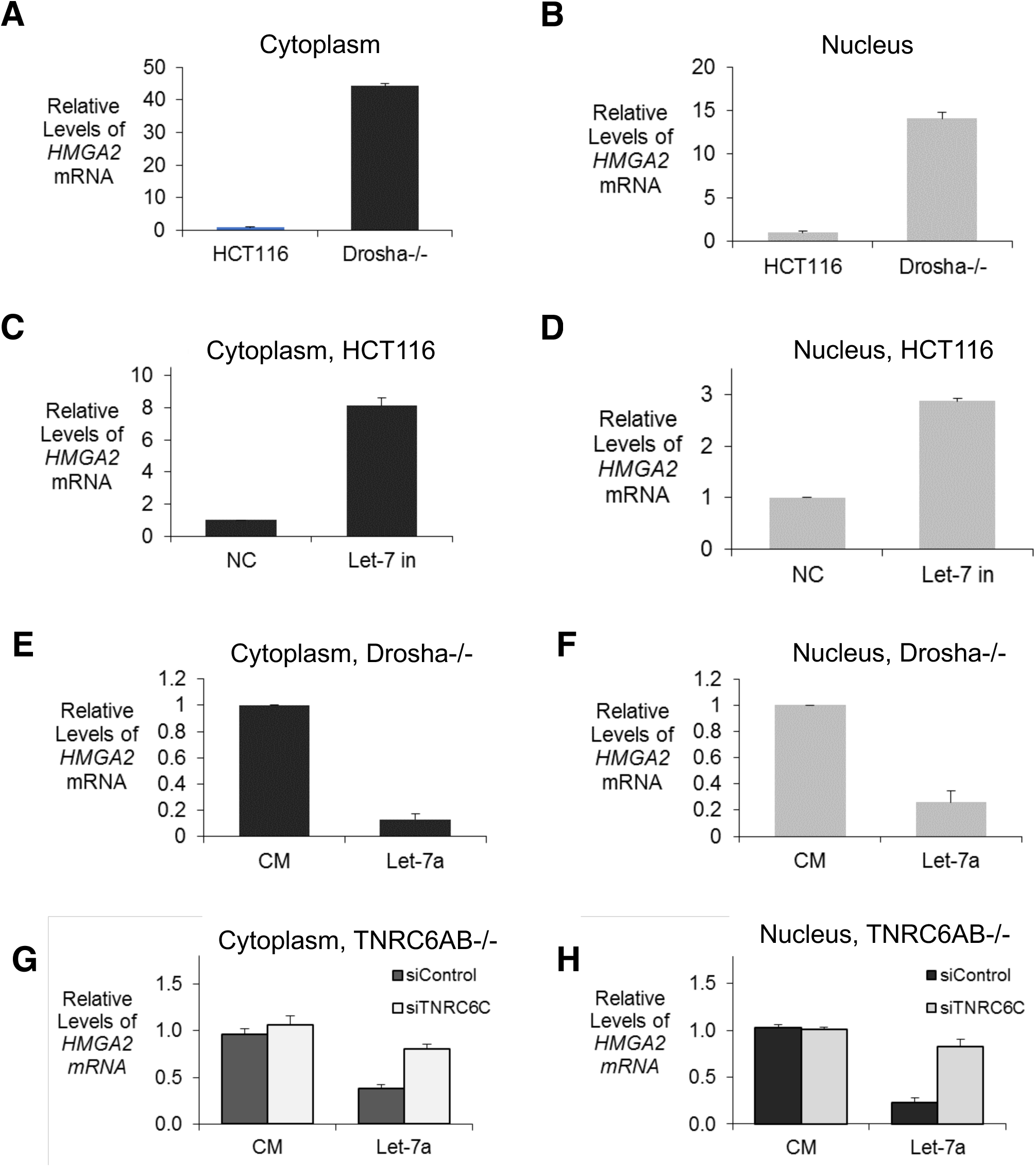
miRNA mediated repression of HMGA2 can occur in the cytoplasm and nucleus. Relative level of mature HMGA2 mRNA in the (A,C,E) cytoplasm and (B,D,F) nucleus of (A,B) WT and Drosha-/-, (C,D) WT cells transfected with *let-7* family miRNA inhibitor and (E,F) Drosha-/- cells transfected with *let-7a* miRNA mimics. Relative level of mature HMGA2 in the (G) cytoplasm and (H) nucleus of TNRC6AB-/- with knockdown of TNRC6C, followed by transfection of non-targeting control or *let-7a* miRNA mimic. All values are plotted as the average of biological replicates +/-SD.

Our examination of the nuclear fraction revealed similar miRNA-mediated regulation of gene expression. In the nuclear fraction of *DROSHA*-/- the expression of *HMGA2* was increased relative to WT (**Figure 5B**). Transfection of *let-7* family miRNA inhibitors led to an increase in HMGA2 expression (**Figure 5D**) while transfection of a *let-7a* miRNA mimic in *DROSHA*-/- cells led to a decrease in expression **(Figure 5F**). Further, silencing of HMGA2 in the nucleus was also dependent on the TNRC6 paralogs, like in the cytoplasm (**Figure 5H**). These observations support the hypothesis that binding of AGO2:mRNA complexes within chromatin-associated RNA and nuclear RNA may also contribute to the silencing of HMGA2.

To determine whether *let-7* mediated inhibition of HMGA2 expression in cell nucleic is observed in other cells, we evaluated HT29 and HeLa cells. The expression of HMGA2 in HT29 and HeLa cells is lower than in HCT116 cells (**Supplementary Figure 7A**). Despite the difference in expression levels, *let-7* mediated regulation was similar. Addition of *let-7* family miRNA inhibitors led to an increase in HMGA2 expression in whole cell (**Supplementary Figure 7B,D**), cytoplasm, and nucleus **(Supplementary Figure 7C,E**).

### HMGA2 expression is not controlled at the level of splicing or transcription

Previous studies from our laboratory and others had demonstrated that synthetic duplex RNAs could control gene transcription and gene splicing (13–28). These previous regulatory duplex RNAs did not target the 3’-UTR. For regulating transcription, the duplex RNAs were complementary to noncoding RNAs that overlap the 5’ or 3’ terminus of target mRNAs (14, 81–83). For gene splicing, they were targeted near intron/exon junctions and splicing regulatory elements (19–21). While the regulatory region within HMGA2 3’-UTR is different from these previous studies, the precedent for nuclear RNAi regulation of transcription or splicing led us to explore the potential for changes in HMGA2 splicing and transcription.

To test the hypothesis that *let-7* miRNAs may regulate the *HMGA2* transcript through modulation of transcription, we first examined the pre-mRNA transcript expression levels. In DROSHA-/- cells, we observed an increase in the *HMGA2* pre-mRNA level **(Figure 6A, Supplementary Figure 9AB)**. Following transfection of *let-7* miRNA family inhibitors in WT **(Figure 6B, Supplementary Figure 9CD)** or *let-7a* miRNA mimics in DROSHA-/- **(Figure 6C, Supplementary Figure 9EF**), we observed no significant change in the level of *HMGA2* pre-mRNA level. Next, we performed RNAPII ChIP-qPCR to measure the recruitment of RNA polymerase II to the HMGA2 promoter. Relative to WT, we observed no change in the recruitment of RNAPII to the HMGA2 promoter in the DROSHA-/- cells **(Figure 6D)**. These data are consistent with the conclusion that indirect effects of *let-7* miRNAs may increase HMGA2 expression, but do not suggest that *let-7* binding to chromatin-associated RNA has a significant impact on HMGA2 transcription.

**Figure 6.**
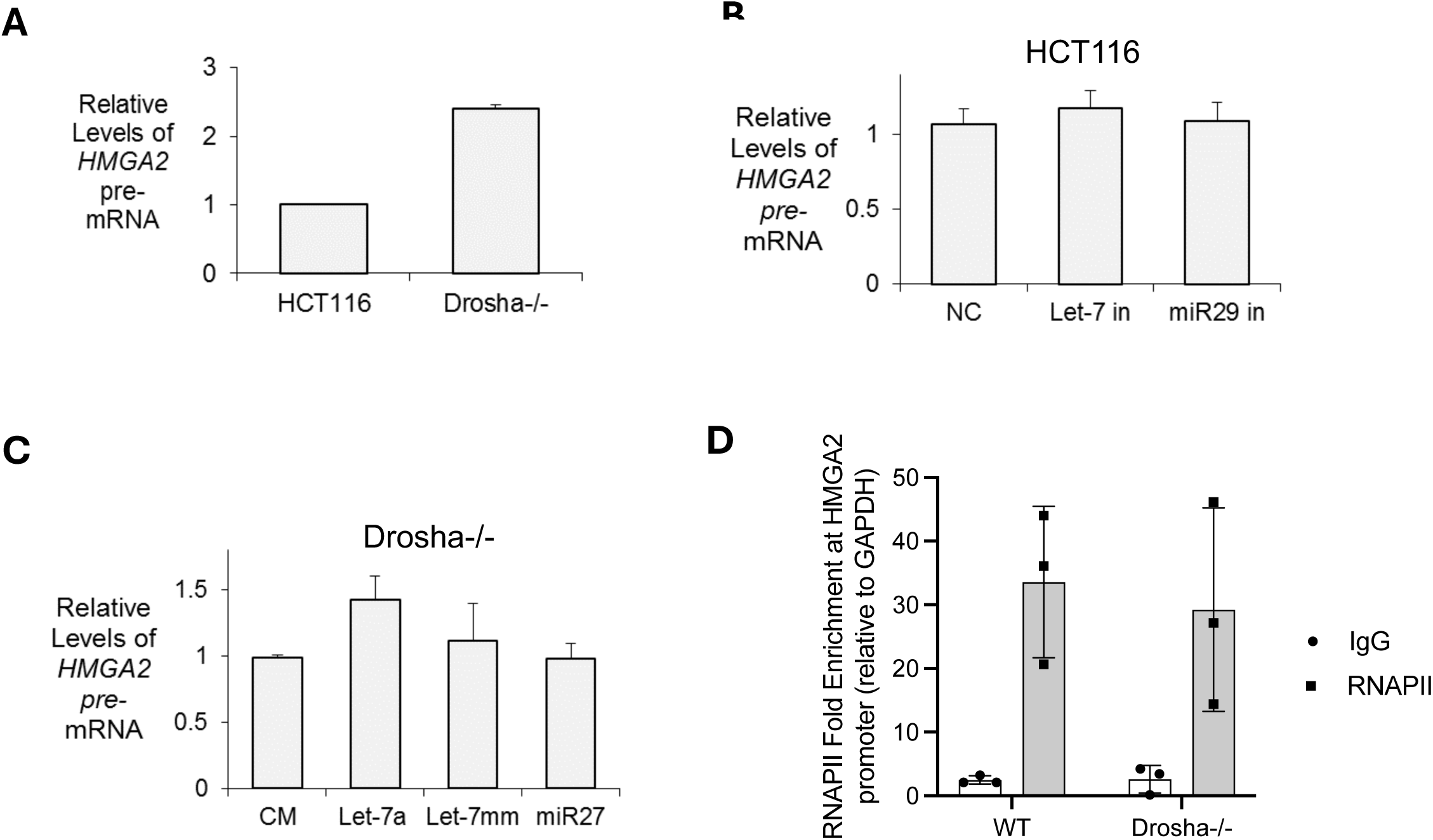
HMGA2 is not transcriptionally regulated by *let-7* family miRNAs. Relative HMGA2 pre-mRNA levels in (A) WT and Drosha-/- cells, (B) WT cells treated with non-targeting miRNA inhibitor control, *let-7* family or *miR-29* family miRNA inhibitor, and (C) Drosha-/- cells transfected with *let-7a* or control miRNA mimics. (D) ChIP-qPCR measuring the recruitment of RNA polymerase II to the HMGA2 promoter in WT and Drosha-/- cells. All values are plotted as the average of biological replicates +/-SD.

Alternatively, a change in HMGA2 expression could also be driven by changes in pre-mRNA splicing. To test this hypothesis, we designed a locus-wide qPCR splicing assay to measure how often a given *HMGA2* exon is included in spliced mRNA compared to how often it is skipped (**Figure 7A**). Primers designed to span exon-exon junctions allow for the specific detection of mRNA isoforms that include or skip a given *HMGA2* exon (**Supplementary Table 1**). Each exon’s inclusion:skipping ratio, relative to the expression of the full-length mRNA, is therefore representative of an exon’s splicing efficiency.

**Figure 7.**
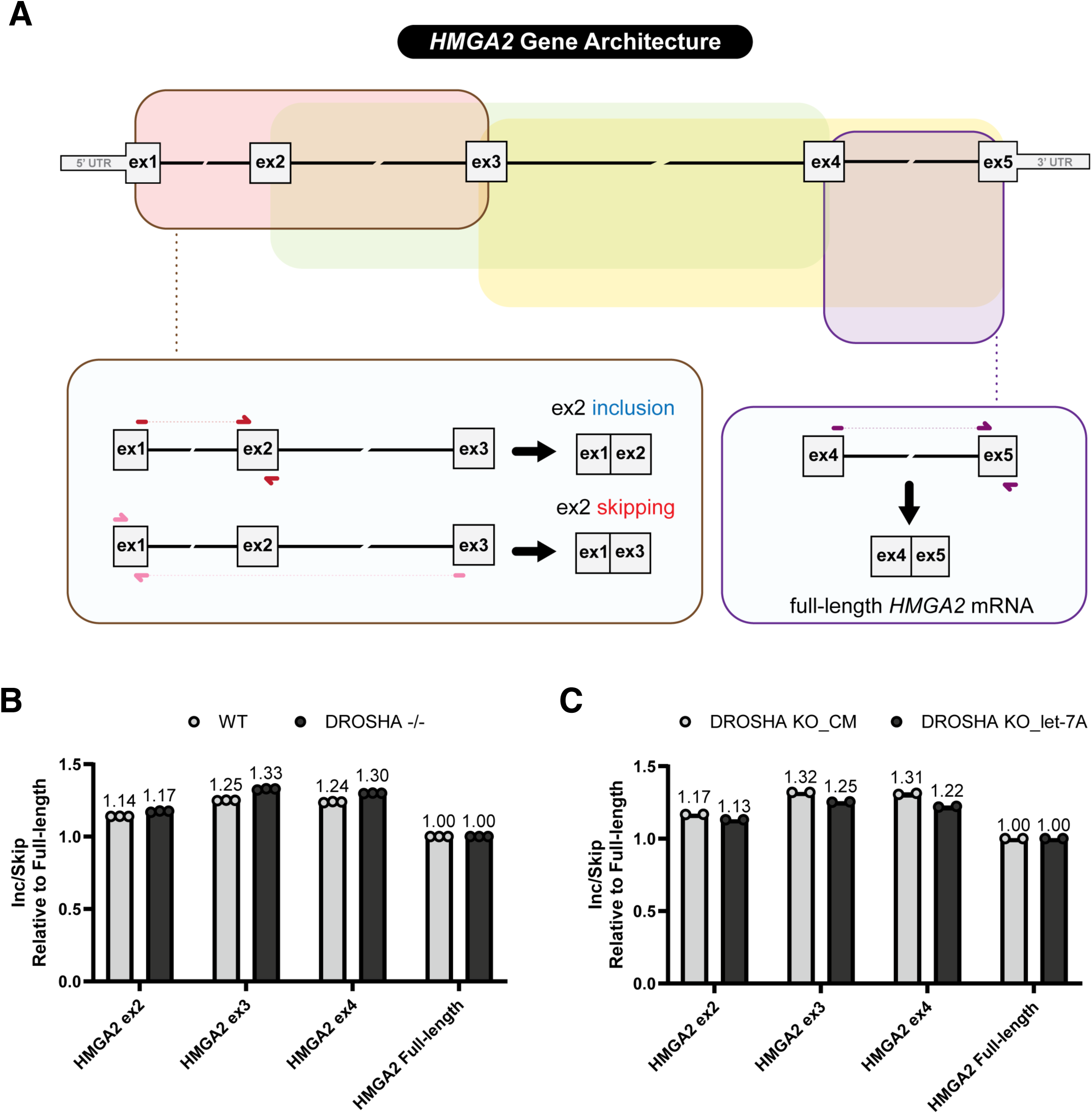
pre-mRNA splicing of HMGA2 is not regulated by *let-7* family miRNAs. **A**) A schematic depicting the exon-exon junction spanning primers designed for the measurement of exon inclusion and/or skipping events within the *HMGA2* gene. qPCR primer pairs were designed to assay splicing of the entire *HMGA2* locus, where each of the four regions targeted are indicated in red, green, yellow, and purple, respectively. Each region has a primer pair directly measuring exon inclusion and another pair measuring exon skipping. With the first region (red) as an example: the presence of solid-colored lines indicate direct complementarity of primers to its target sequence, whereas dashed-colored lines indicate no complementarity. The results from this assay are plotted as splicing inclusion:skipping ratios, as shown in B) for untreated WT and DROSHA-/- and C) for DROSHA-/- transfected with let-7a miRNA mimic or control. Each condition’s respective mean value, calculated from independent biological replicates, are displayed above each condition shown.

We first applied this assay to understand how loss of mature miRNAs in the *DROSHA*-/- cells might affect *HMGA2* gene splicing. Relative to WT HCT116 cells, we observed that the splicing efficiency of each *HMGA2* exon is similar in the DROSHA-/- and wild-type cells (**Figure 7B**). To further test the hypothesis that *let-7* miRNAs may regulate *HMGA2* gene splicing, we repeated the same splicing assay using *DROSHA*-/- cells transfected with either a *let-7a* mimic or a non-complementary (CM) control. We observed only minor differences in splicing efficiency relative to the CM control (**Figure 7C**). A similar qPCR splicing assay done in parallel targeting the reference gene *TBP*, an internal control for each experimental condition presented, demonstrates that miRNA-mediated splicing modulation of *HMGA2* by *let-7a* is sequence specific (**Supplementary Figure 10**). Our data do not support the conclusion that *let-7* binding has a major effect on splicing within *HMGA2* mRNA.

## Discussion

### Cytoplasmic and nuclear RNAi

Complexes between AGO and RNA are programmable recognition elements. Ago protects the RNA and facilitates target recognition while, the small RNA component directs the complex to target sequences. While the canonical targets for miRNAs are within the 3’-UTRs of cytoplasmic RNAs (1–3), there is no theoretical reason why sites outside the 3’-UTR cannot be targeted, including targets within mammalian cell nuclei.

Synthetic duplex RNAs can complex with AGO to control gene transcription (13–18) and regulate gene splicing (19–21). AGO:RNA complexes complementary to the disease-associated repeat RNAs of Friedreich’s Ataxia (84, 85), Fuchs Dystrophy (86, 87), and C9orf72-related ALS (88) can block R-loop formation or protein association. Localization of AGO protein to cell nuclei when cells are grown to high confluence can de-repress cytoplasmic miRNA targets (50). While these reports are intriguing, most work has focused on the action of synthetic, designed RNAs. The unanswered question is whether the robust impacts of designed duplex RNAs in cell nuclei are reflected in widespread, biologically significant effects on the regulation of cellar gene expression by miRNAs. Our results have identified the potential for RISC to bind chromatin-associated RNA and silence endogenous nuclear targets, suggesting a degree of nuclear control of gene expression by miRNAs and RNAi factors.

### Identification *HMGA2:let-7* regulation using anti-AGO2 chimeric eCLIP

Identifying suspected interactions between miRNAs and RNA targets can be accomplished with computational analysis to locate seed sequence complementarity. Computational analysis, however, will identify many potential sites for biological regulation. The challenge is to identify methods that prioritize sites that are most likely to be significant for cellular regulation.

To meet our criteria as a “high priority target”, a site was required to meet the following criteria: 1) non-chimeric AGO2 peak density; 2) chimeric AGO2 peak density; 3) the miRNAs revealed by chimeric eCLIP belong to well-expressed miRNA families; and 4) there be a significant change in the expression of target genes in wild type versus *DROSHA*-/- cells.

Our chimeric eCLIP data revealed few genes that met these criteria (**Supplementary Table 2**). Further experimental validation showed that only one of these candidates (**Figure 3-5, Supplementary Figures 4-7**), *HMGA2*, was a strong candidate for direct repression by miRNAs in HCT116 cells. Regulation by *let-*7 miRNAs was supported by data showing increased expression upon addition of *let-7* family miRNA inhibitors (**Figure 4B**), rescued gene silencing in DROSHA-/- cells following *let-7* miRNA mimic transfection (**Figure 4C**), and modulation in HMGA2 expression by inhibitors and mimics following knockout and knockdown of the TNRC6 paralogs (**Figure 4D**).

While *let-7* had been characterized as a regulator of *HMGA2* in other cell types (66–69), it is a good example of the power of chimeric eCLIP for prioritizing candidates. A PubMed search of “miRNA” and HCT116 reveals over 1300 publications. Chimeric eCLIP identified HMGA2 as a priority candidate for experimental validation in HCT116 cells, even though many reports had implicated other miRNAs and other gene targets. Several of these publications implicate *HMGA2* as a target, but to our knowledge none discuss *let-7* as partner. Other publications discuss *let-7*, but do not focus on H*MGA2* as a target. Using chimeric eCLIP, however, we identified non-chimeric AGO2 association and chimeric binding peaks in the HMGA2 3’UTR in multiple subcellular fractions. These chimeric binding interactions were predominately found to involve *let-7* family miRNAs, one of the most abundant miRNA families in HCT116 cells. Our experimental confirmed the priority assigned to *HMGA2* by the chimeric eCLIP data.

### Nuclear Recognition and Regulation

Our replicate chimeric eCLIP data using chromatin from wild-type HCT116 cells reveals binding of AGO2 and chromatin-associated RNA throughout the transcriptome. While most nonchimeric reads are within introns, most chimeric reads and the most promising candidate interactions are within 3’-UTRs. Our data from nuclear lysate using HCT116-NLS cells where AGO2 is almost exclusively nuclear show similar distribution of binding for miRNA:AGO2 complexes. Our replicate data from HCT116 cytoplasm is also similar.

Our initial data identified *HMGA2* and a handful of other genes as prime candidates for miRNA-mediated recognition of chromatin-associated RNA. Experimental validation revealed that nuclear *HMGA2* is regulated directly by miRNAs in HCT116 cells (**Figure 5**). Recognition begins when *HMGA2* mRNA is associated with chromatin (**Figure 3**), but this binding does not affect transcription or splicing (**Figures 6 and 7**).

While beyond the scope of this report, we are investigating the molecular mechanism of miRNA-mediated regulation of *HMGA2* expression in cell nuclei. It is possible that the mechanism shares similarities with the known cytoplasmic mechanism for gene silencing. Previous work from our laboratory and others have demonstrated the association of members of the CCR4-NOT complex with AGO2 and TNRC6A in the nucleus by mass spectrometry and subsequently confirmed these associations by reciprocal immunoprecipitations of individual proteins (16, 56). The similarity of protein complexes supports the hypothesis that the mechanisms of regulation will also be similar and that both mechanisms will contribute to the efficiency of miRNA-mediated gene silencing.

### Strengths and limitations of chimeric eCLIP

These data demonstrate that chimeric eCLIP is a valuable approach for rapidly and unambiguously identifying miRNA:RNA target interactions that may be biologically relevant. We found that chimeric eCLIP was reproducible - we obtained the similar data from replicate chimeric eCLIP experiments. The complementary chimeric and nonchimeric data affords high confidence that the data reflects cellular interactions that may be biologically relevant. Taken together our experience with whole cell and chromatin-bound RNA samples demonstrates that chimeric eCLIP is an excellent experimental starting point for identifying miRNA:target interactions. The ability of chimeric eCLIP to provide data necessary for decisive candidate identification is consistent with it being a powerful approach to finding some of the most powerful potential regulatory miRNA/target pairs in different cell types or tissues.

While exciting, it is necessary to recognize that chimeric eCLIP protocols will need to be improved to reach their full potential. Like all methods, chimeric eCLIP has limitations. One limitation more reflects the complexity of miRNA regulation than a shortcoming of the technology. Strong chimeric eCLIP data does not prove the existence of miRNA-mediated regulation. For example, we observe strong evidence of AGO2 and *let-7* binding at the *MYC* gene but obtained no experimental evidence supporting miRNA-mediated gene regulation (**Supplementary Figure 4**). It is likely that, for some genes, the sensitivity of eCLIP data will reveal a real association between miRNAs and RNA targets. These associations are efficient enough to produce signal that can be detected by chimeric eCLIP but that are not efficient enough to substantially repress expression of the target genes. That hypothesis may explain why we observe strong AGO2:*let7* association within the *MYC* 3’-UTR but were unable to obtain experimental evidence for on-target regulation of *MYC* expression by *let-7*.

Another limitation reflects the boundaries of the current technology and the fact that, given equally efficient binding of miRNAs, well-expressed targets will be detected better than targets that are less well expressed. Transcripts that are not well expressed, are unstable, or are not accessible may not be detected with confidence or may not be detected at all. Compounding this shortcoming, the need for a ligation step makes detection of chimeric reads inefficient, further emphasizing recognition of well-expressed transcripts. Rather than detecting millions of reads as is common with standard anti-AGO2 eCLIP, chimeric eCLIP detects only tens of thousands of chimeric reads and may miss biologically important interactions. The practical consequence of this limitation for our interest in nuclear RNAi is that we may be missing target sequences within intronic RNA that affect splicing or targets within promoter RNAs that affect transcription.

Technical innovation will be necessary to improve chimeric eCLIP and overcome these limitations. Methods are being developed that focus the power of chimeric eCLIP on specific genes, allowing much deeper insights at loci when miRNA-mediated control is suspected (84). These methods may be especially important for investigating relatively lowly abundant sequences associated with chromatin, especially intronic sequences. While our chimeric eCLIP approach was unbiased and eventually identified sequences with 3’-UTRs as primary miRNA targets, we began the project to test the hypothesis that we would identify nuclear RNA targets for the regulation of splicing, transcription, or other processes. Our analysis at the *HMGA2* locus did not identify did not implicate regulation of transcription or splicing.

More broadly, our chimeric eCLIP did not reveal strong association within intronic or other noncoding RNAs. As noted above, one explanation is that these noncoding targets may be expressed at low levels too low for identification by chimeric eCLIP. Improved methods for targeted chimeric eCLIP (48) may alleviate this problem. Our nonchimeric data (Tse unpublished), however, does reveal some potential sites for interactions between AGO2 and introns and we are investigating these as candidate sites that may contribute to the regulation of splicing. Alternatively, as we and others have shown, miRNAs and RNAi factors can shift within the cell depending on environmental conditions (22, 27, 50, 89–91). miRNA-mediated regulation may become both observable and biologically important under a special set of cell growth of physiological conditions, and sensitive methods like chimeric eCLIP would need to be applied to those conditions.

### Conclusions

Our data suggest that AGO2:miRNA complexes associate with chromatin in mammalian cells. More broadly, chimeric eCLIP is a powerful method for prioritizing gene candidates for miRNA-mediated regulation that merits wide use in studies that aim to efficiently identify miRNA:RNA target pairings in cells and tissue. The most obvious associations identified by chimeric eCLIP in chromatin or nucleoplasm occur within the 3’-UTR and binding sites are like those observed in the cytoplasm. We have validated *HMGA2* as target for *let-7* in both the nucleus and cytoplasm, with RNA levels reduced in both compartments. Future studies will be needed that focus on the molecular mechanism of this miRNA-mediated gene repression in cell nuclei.

Our data demonstrates that miRNAs can recognize sequences within chromatin-associated RNAs and suggests that recognition within the 3’-UTR can contribute to miRNA-mediated control of gene expression. While previous studies have shown that nuclear RNAs can regulate splicing or transcription and we observe recognition of chromatin-associated RNA, we do not observe miRNA mediated control of either process at the *HMGA2* loci. While powerful, improved protocols for chimeric eCLIP or similarly innovative methods may be necessary to better test the hypothesis that biologically active miRNAs can function by targeting intronic, promoter, or other less expressed RNAs.

Using chimeric eCLIP, we identified *HMGA2*, a benchmark gene that allowed us to probe miRNA-mediated regulation in HCT116 cytoplasm and nuclei. More broadly, our experience with chimeric eCLIP paints a promising picture of its broad potential to identify miRNA:RNA target interactions. The data was robust and internally consistent, providing confident identification of major sites for binding by AGO2:miRNA complexes that justifies the time-consuming experimental validation necessary to link miRNA recognition to on-target regulation of a specific gene. In this case, the identification of nuclear and cytoplasmic RNA targets for miRNA and how recognition in both compartments cooperate to shape gene regulation may impact understanding of human cell biology and drug discovery.

## Acknowledgements

This study was supported by R35GM118103 (DRC) from the National Institutes of Health and the Welch Foundation (I-2184 to DRC). DRC is the Rusty Kelley Professor of Biomedical Science. We thank our colleagues from EclipseBio for providing text describing chimeric eCLIP methodology

## Author Contributions

D.R.C. helped write the manuscript, design experiments, and analyze data. C.H., J.H. R.B, and V.T. designed experiments, performed experiments, analyzed experiments, and helped write the manuscript.

## SUPPLEMENTARY INFORMATION

**Supplementary table 1.**
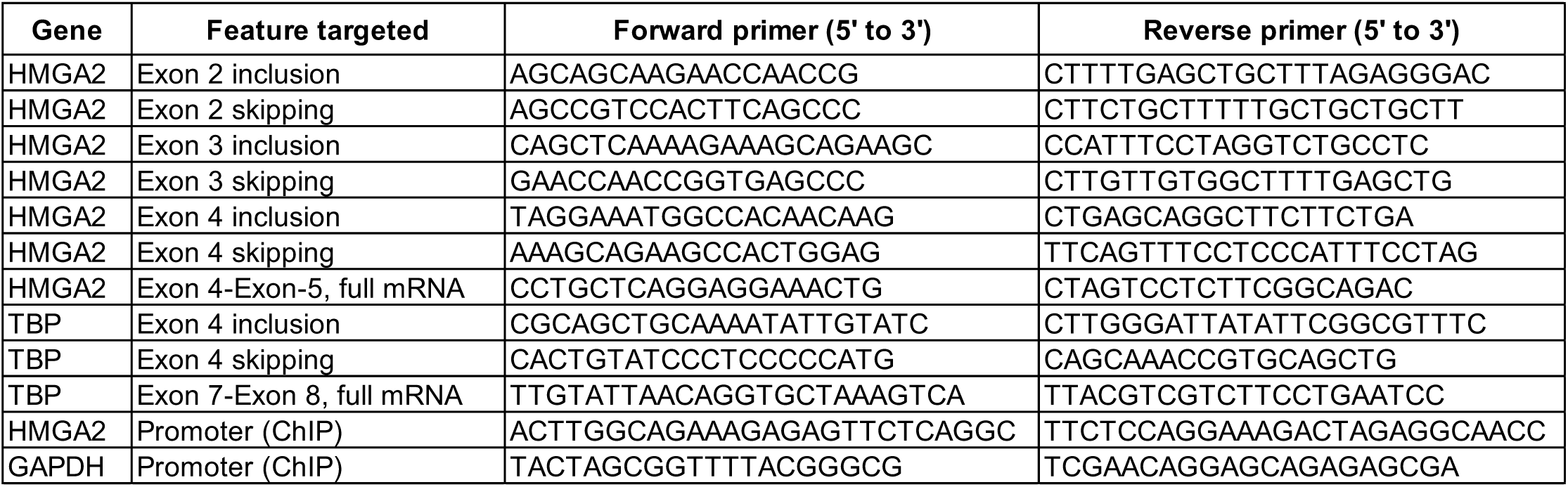
All qPCR primers used in splicing, ChIP and RT-qPCR assays

**Supplementary table 2.** Argonaute 2 chimeric binding sites in gene 3’UTR identified in the (A) cytoplasm, (B) nucleus and (C) chromatin. Genes were identified based on number of chimeric binding peaks per gene and conservation of binding peaks across subcellular fractions. Peaks were defined to have a p-value<0.001 and log2FC>3 (IP vs. input).

**A**
| Cytoplasm |  |  |  |  |  |
| --- | --- | --- | --- | --- | --- |
| Gene | Strand | Gene expression FC (Drosha<sup>-/-</sup> vs. WT) | Ago2 chimeric peaks with miRNA seed match in 3' UTR | Number of unique miRNAs | All miRNAs |
| HMGA2 | + | 5.0 | 26 | 15 | miR-20a-5p, miR-93-5p, miR-106b-5p, miR-29a-3p, let-7a-5p, let-7d-5p, let-7f-5p, miR-17-5p, let-7b-5p, let-7g-5p, miR-16-5p, miR-33b-5p, miR-15a-5p, miR-15b-5p |
| MYC | + | -8.8 | 7 | 5 | let-7a-5p, let-7b-5p, let-7d-5p, let-7e-5p, miR-320a-3p |
| TNFRSF12A | + | 1.4 | 2 | 2 | miR-19b-3p, miR-19a-3p |
| ZFP36 | + | -2.3 | 6 | 6 | miR-222-3p, miR-31-5p, miR-27a-3p, miR-200c-3p, miR-29b-3p, miR-29a-3p |
| ZFP36L1 | - | 3.4 | 8 | 7 | miR-222-3p, let-7a-5p, miR-23a-3p, miR-31-5p, miR-18a-5p, miR-29b-3p, miR-29a-3p |

**B**
| Nucleus |  |  |  |  |  |
| --- | --- | --- | --- | --- | --- |
| Gene | Strand | Gene expression FC (Drosha<sup>-/-</sup> vs. WT) | Ago2 chimeric peaks with miRNA seed match in 3' UTR | Number of unique miRNAs | All miRNAs |
| HMGA2 | + | 5.0 | 8 | 8 | let-7a-5p, let-7d-5p, miR-106b-5p, miR-20a-5p, miR-17-5p, let-7b-5p, let-7f-5p, miR-16-5p |
| MYC | + | -8.8 | 2 | 2 | let-7b-5p, let-7d-5p |
| TNFRSF12A | + | 1.4 | 2 | 2 | miR-19b-3p, miR-19a-3p |
| ZFP36 | + | -2.3 | 2 | 2 | miR-31-5p, miR-27a-3p |
| ZFP36L1 | - | 3.4 | 1 | 1 | miR-29b-3p |

**C**
| Chromatin |  |  |  |  |  |
| --- | --- | --- | --- | --- | --- |
| Gene | Strand | Gene expression FC (Drosha<sup>-/-</sup> vs. WT) | Ago2 chimeric peaks with miRNA seed match in 3' UTR | Number of unique miRNAs | All miRNAs |
| HMGA2 | + | 5.0 | 8 | 5 | miR-16-5p, let-7d-5p, let-7a-5p, let-7f-5p, let-7b-5p |
| MYC | + | -8.8 | 1 | 1 | let-7b-5p |
| TNFRSF12A | + | 1.4 | 2 | 2 | miR-19b-3p, miR-19a-3p |

**Supplementary figure 1.**
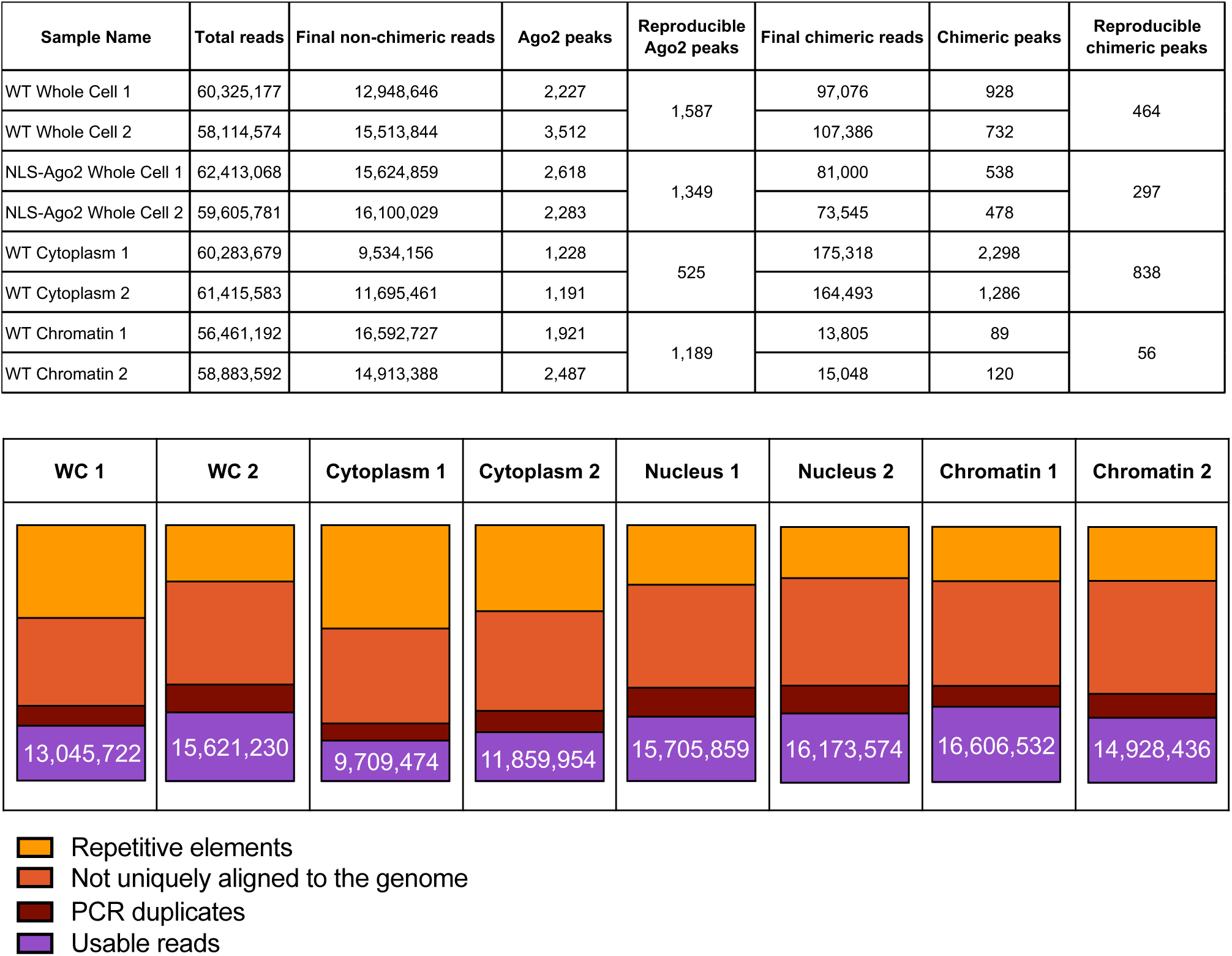
Ago2 chimeric eCLIP quality control. (A) Number of reads (total, non-chimeric and chimeric), AGO2 peaks (non-chimeric and chimeric), and reproducible AGO2 peaks (non-chimeric and chimeric) for all collected sample types analyzed by chimeric eCLIP. (B) Total reads from each replicate of chimeric eCLIP aligning as repetitive elements, not uniquely aligning to the genome, PCR duplicates, and usable reads.

**Supplementary figure 2.**
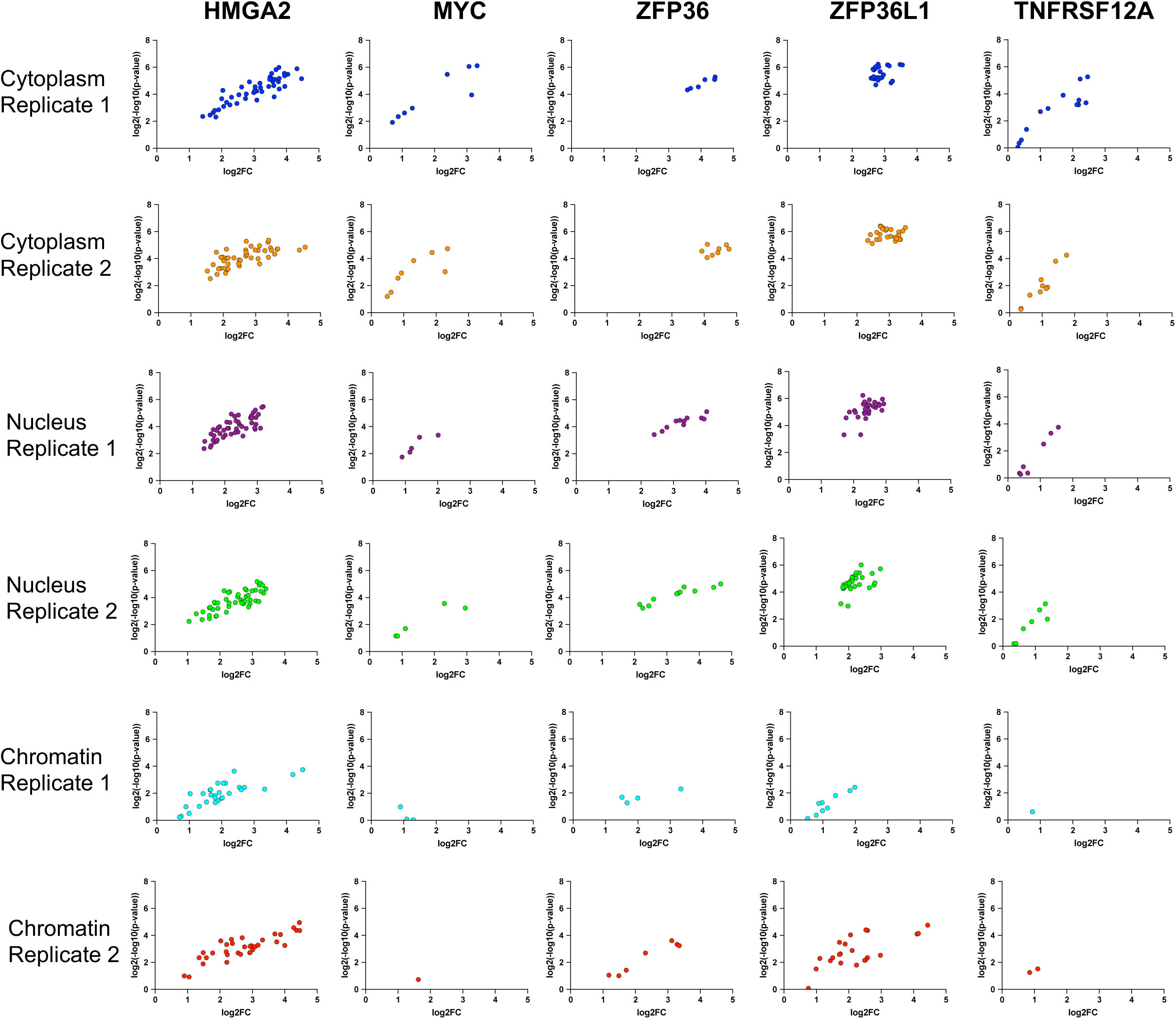
Strength of individual non-chimeric AGO2 peaks in the 3’UTR of genes from supplemental table 1. All non-chimeric AGO2 peaks in the 3’UTR of the five genes (from Supplemental Table 2) are plotted as log2FC (IP vs. input) vs. peak significance (log2(-log10(p-value)), with each dot representing a single AGO2 peak. Following IP and sequencing, peaks were identified following read alignment to the genome and identification using CLIPper.

**Supplementary Figure 3.**
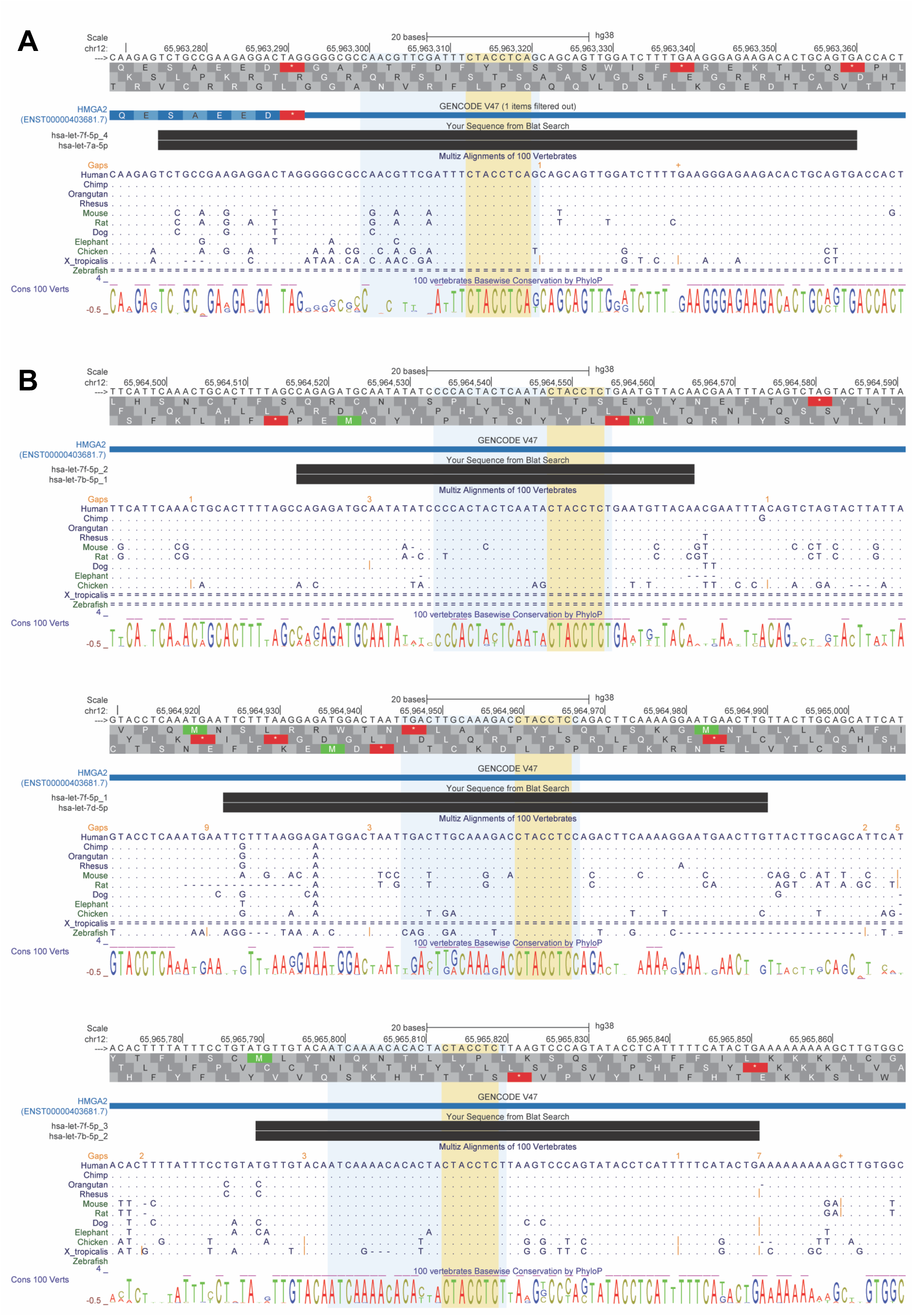
The seed sequence of Let-7 miRNAs from this study are strongly conserved across species within the 3′ UTR of *HMGA2*. **(A)** Alignment and comparison of let-7 miRNAs with perfect matches to seed sequences. **(B)** Alignment and comparison of let-7 miRNAs with imperfect matches to seed sequences. The genomic coordinates, followed by the reading frames for canonical and alternative amino acid sequences (top and bottom, respectively), including all nucleotides shown in (A) and (B) for *HMGA2*, NM_003483.6 (ENSG00000149948.14), corresponds to the sense strand of the gene. The span of the entire miRNA sequence is indicated in light blue whereas the seed sequence is indicated in yellow. The canonical full-length transcript for *HMGA2* of the human genome is indicated in blue under the GENCODE v47 track. The 3′ UTR is indicated by a thin box whereas a thicker box with amino acid symbols corresponds to the coding sequence. CLEAR-CLIP sequence contexts aligned to the human genome is indicated in black boxes under the BLAT Search track. Multiple alignment and comparison of genomes are indicated by their species under the Multiz Alignments of 100 Vertebrates track, in addition to their basewise conservation across species under the 100 Vertebrates Basewise Conservation by PhlyoP track. Under the Multiz Alignment track, each dot for a given species at a given position represents an identical nucleotide match to the human reference genome. Each single or double line for a given species at a given position corresponds to no nucleotide(s) or unalignable nucleotides to the reference, respectively. Gold line and text for a given species corresponds to gaps in the region of “N” nucleotide length (a gap of “+” indicates a larger than mappable gap on the UCSC Genome Browser viewer). Vertical blue bars and green square brackets correspond to genomic breaks (e.g., such is the case for Zebrafish; the genomic break occurs at GRCh38/hg38 coordinates chr12:65,965,479-65,965,479, resulting in a proceeding empty row observed in the last snapshot of (B)).

**Supplementary figure 4.**
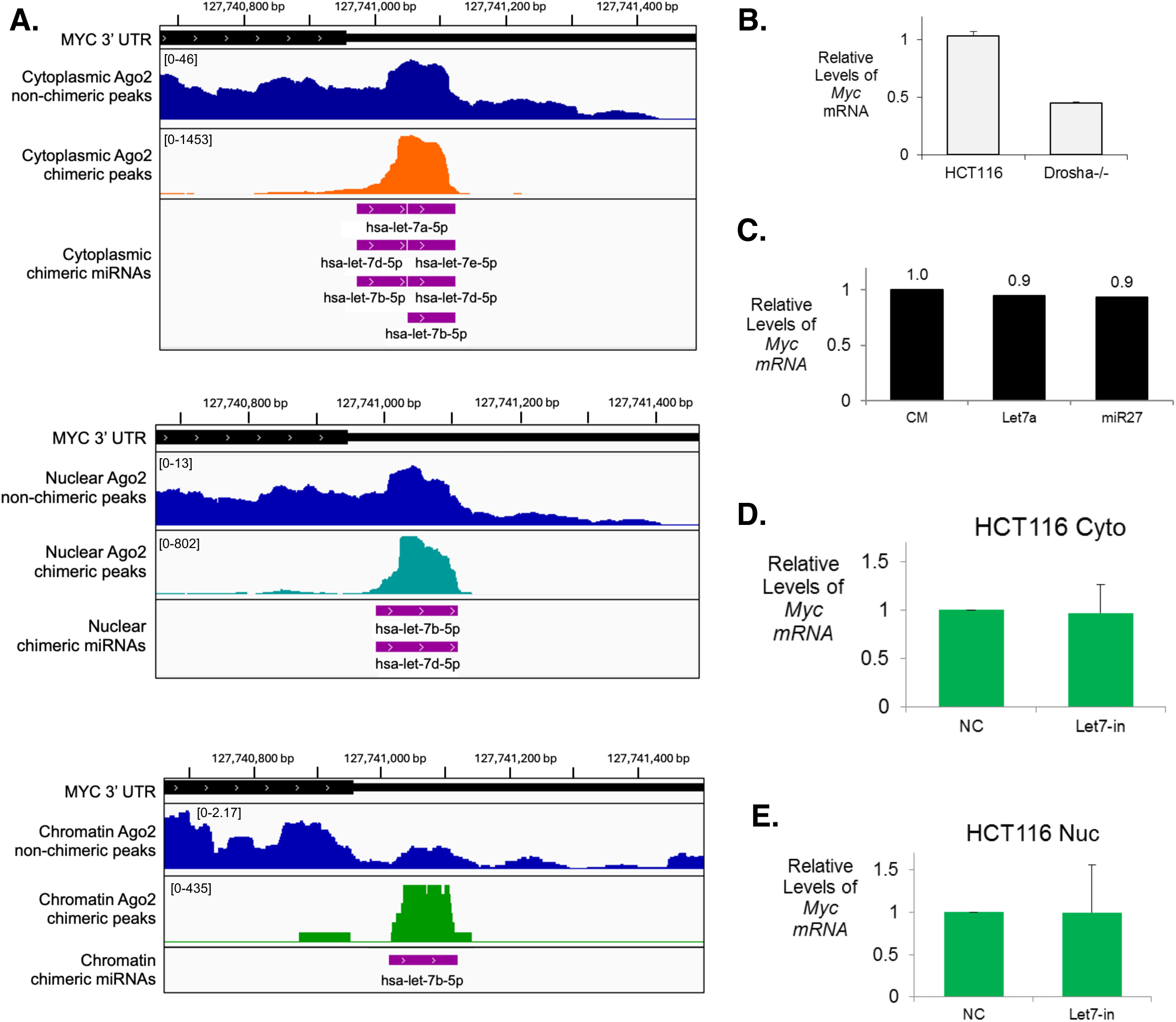
MYC is a miRISC target that is not clearly regulated by the *let-7* family of miRNAs (**A)** Representative IGV browser views of chimeric eCLIP reads within the MYC 3’UTR in the cytoplasm (top), nucleus (middle), and chromatin (bottom). Mature MYC mRNA levels in (**B)** WT and Drosha-/- cells, (**C)** Drosha-/- cells transfected *with let-7a* miRNA mimics, and WT (**D)** cytoplasm and (**E)** nucleus fractions following treatment with *let-7* family miRNA inhibitors. Values are plotted as the average +/- SD.

**Supplementary Figure 5.**
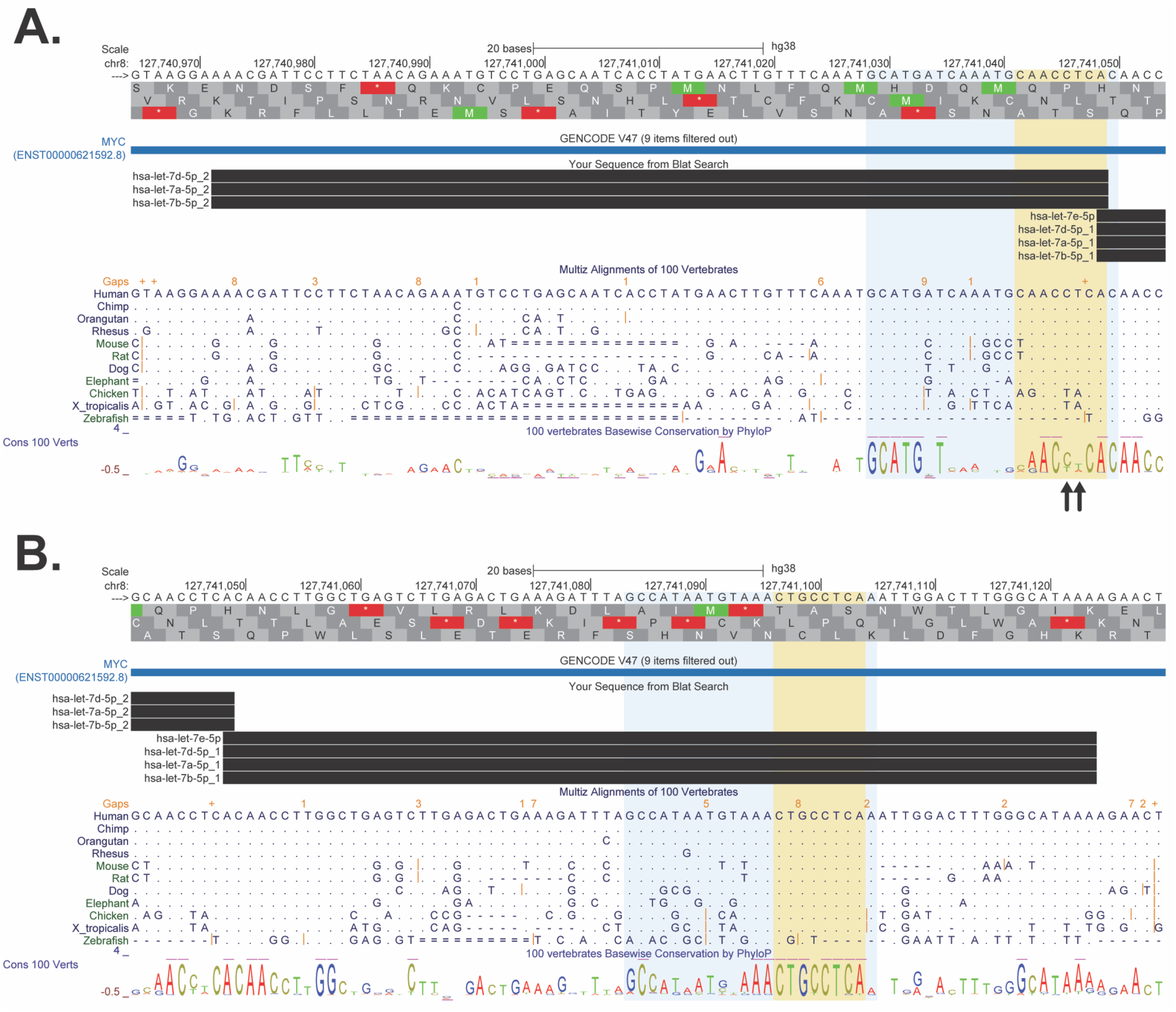
The seed sequence of Let-7 miRNAs from this study are also strongly conserved across species within the 3′ UTR of *MYC*. **(A)** Alignment and comparison of let-7 miRNAs with imperfect matches to seed sequences that exhibit relatively strong conservation. **(B)** Alignment and comparison of let-7 miRNAs with imperfect matches to seed sequences that exhibit strong conservation. The genomic coordinates, followed by the reading frames for canonical and alternative amino acid sequences (top and bottom, respectively), including all nucleotides shown in (A) and (B) for *MYC*, NM_002467.6 (ENSG00000136997.22), corresponds to the sense strand of the gene. The span of the entire miRNA sequence is indicated in light blue whereas the seed sequence is indicated in yellow. The canonical full-length transcript for *MYC* of the human genome is indicated in blue under the GENCODE v47 track. The 3′ UTR is indicated by a thin box whereas a thicker box with amino acid symbols corresponds to the coding sequence. CLEAR-CLIP sequence contexts aligned to the human genome is indicated in black boxes under the BLAT Search track. Multiple alignment and comparison of genomes are indicated by their species under the Multiz Alignments of 100 Vertebrates track, in addition to their basewise conservation across species under the 100 Vertebrates Basewise Conservation by PhlyoP track. Under the Multiz Alignment track, each dot for a given species at a given position represents an identical nucleotide match to the human reference genome. Each single or double line for a given species at a given position corresponds to no nucleotide(s) or unalignable nucleotides to the reference, respectively. Gold line and text for a given species corresponds to gaps in the region of “N” nucleotide length (a gap of “+” indicates a larger than mappable gap on the UCSC Genome Browser viewer). Vertical blue bars and green square brackets correspond to genomic breaks.

**Supplementary figure 6.**
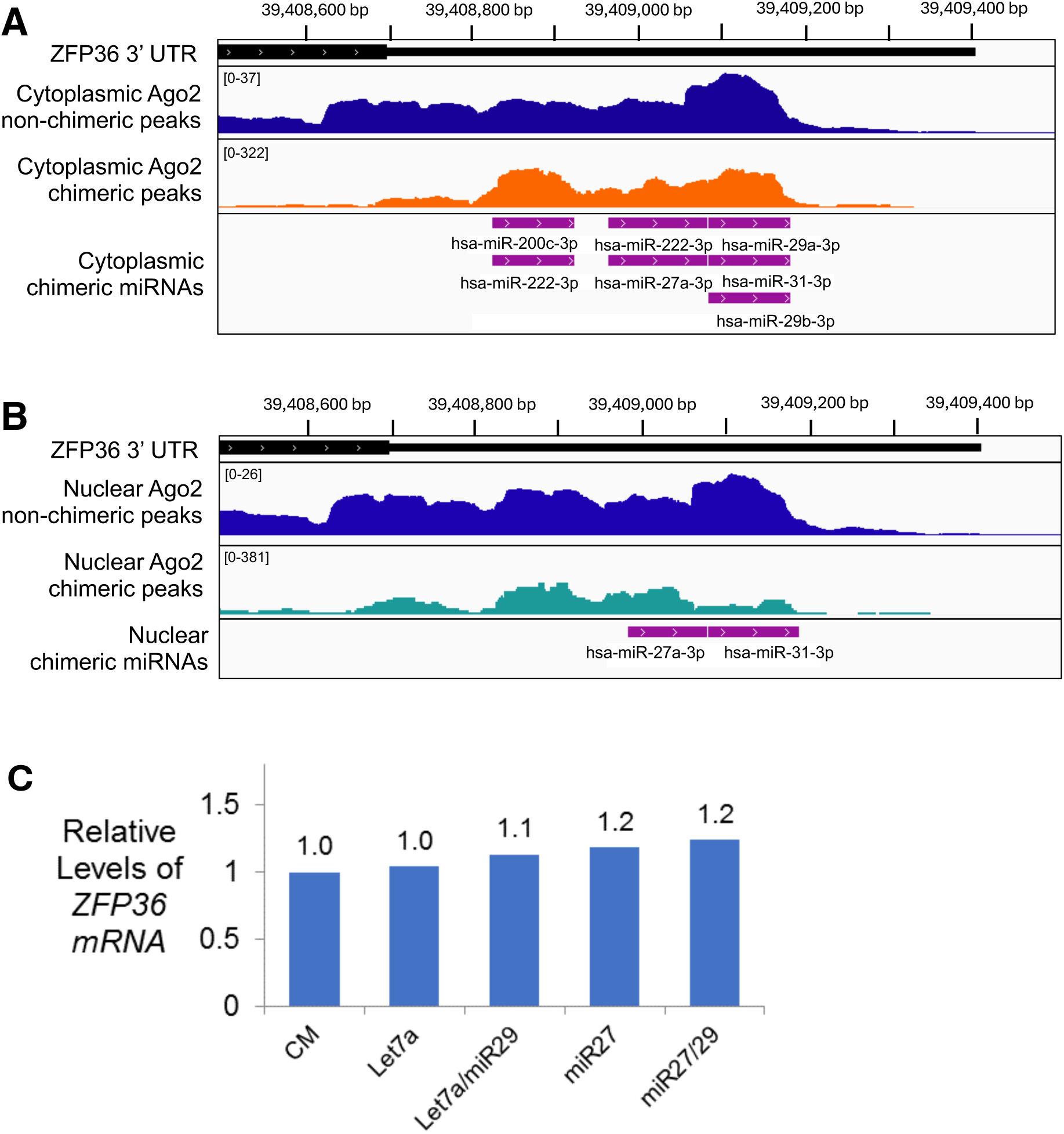
ZFP36 does not seem to be simply regulated by abundant miRNA families. IGV browser views of non-chimeric and chimeric AGO2 binding in the (A) cytoplasm and (B) nucleus. (C) Relative level of mature ZFP36 mRNA in DROSHA-/- cells transfected with miRNA mimics and controls. Values are plotted as the average +/-SD.

**Supplementary figure 7.**
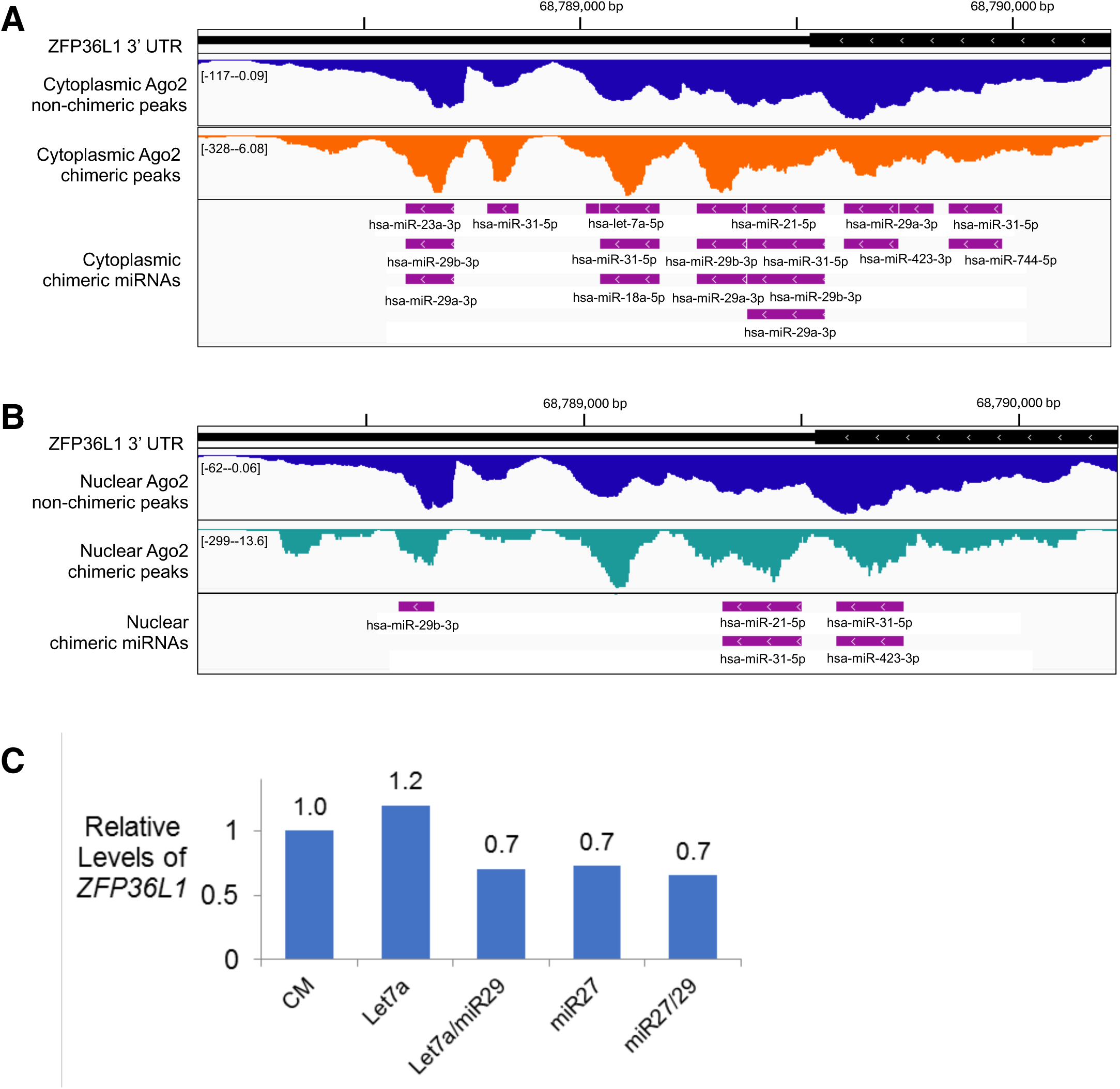
ZFP36L1 does not seem to be simply regulated by abundant miRNA families. IGV browser views of non-chimeric and chimeric AGO2 binding in the (A) cytoplasm and (B) nucleus. (C) Relative level of mature ZFP36L1 mRNA in DROSHA-/- cells transfected with miRNA mimics and controls. Values are plotted as the average +/-SD.

**Supplementary figure 8.**
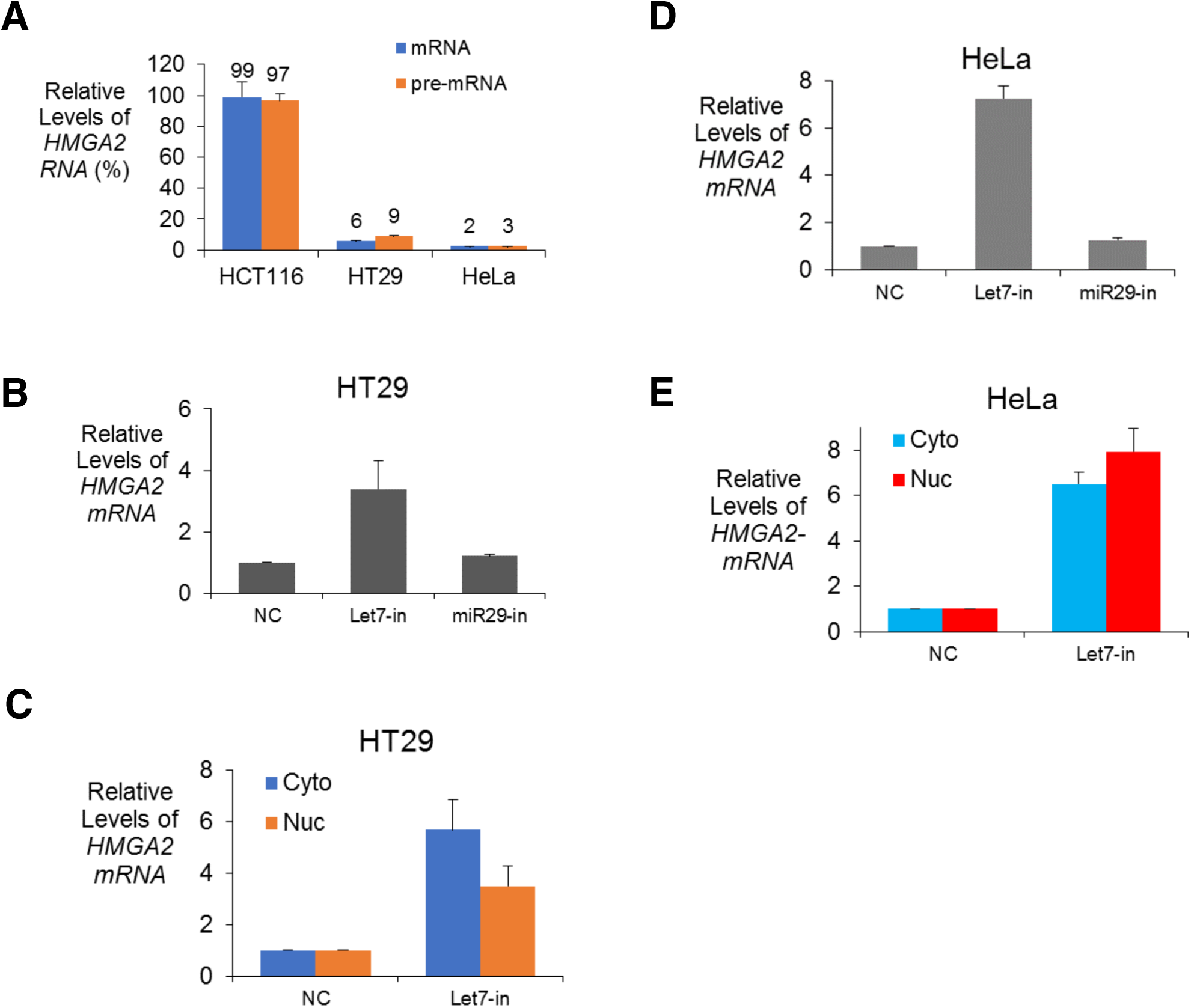
HMGA2 is silenced by *let-7* family miRNAs in other cell lines. (A) Relative level of mature HMGA2 mRNA in HCT116, HT29, and HeLa cells. Relative level of HMGA2 mature mRNA in (B,D) whole cell, (C,E) cytoplasm, and nucleus following transfection of let-7 family miRNA inhibitor and control miRNA inhibitors in (B,C) HT29 and (D,E) HeLa cells. Values are plotted as the average +/-SD.

**Supplementary figure 9.**
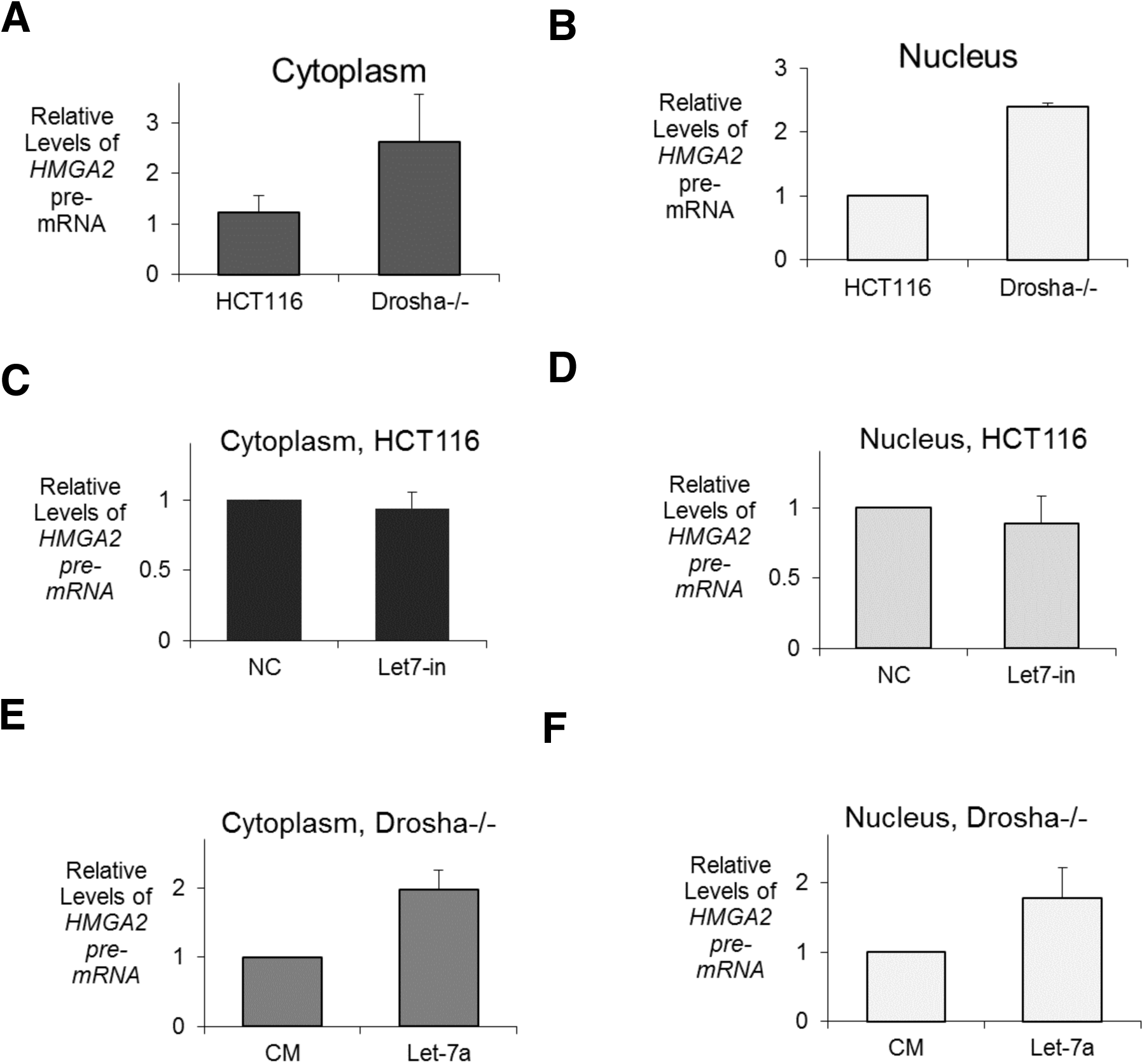
Impact of *let-7* family miRNA on HMGA2 pre-mRNA levels. Relative level of HMGA2 pre-mRNA in the (A,C,E) cytoplasm and (B,D,F) nucleus of (A,B) WT and Drosha-/-, (C,D) WT cells transfected with *let-7* family miRNA inhibitor and (E,F) Drosha-/- cells transfected with *let-7a* miRNA mimics. Values are plotted as the average +/-SD.

**Supplementary figure 10.**
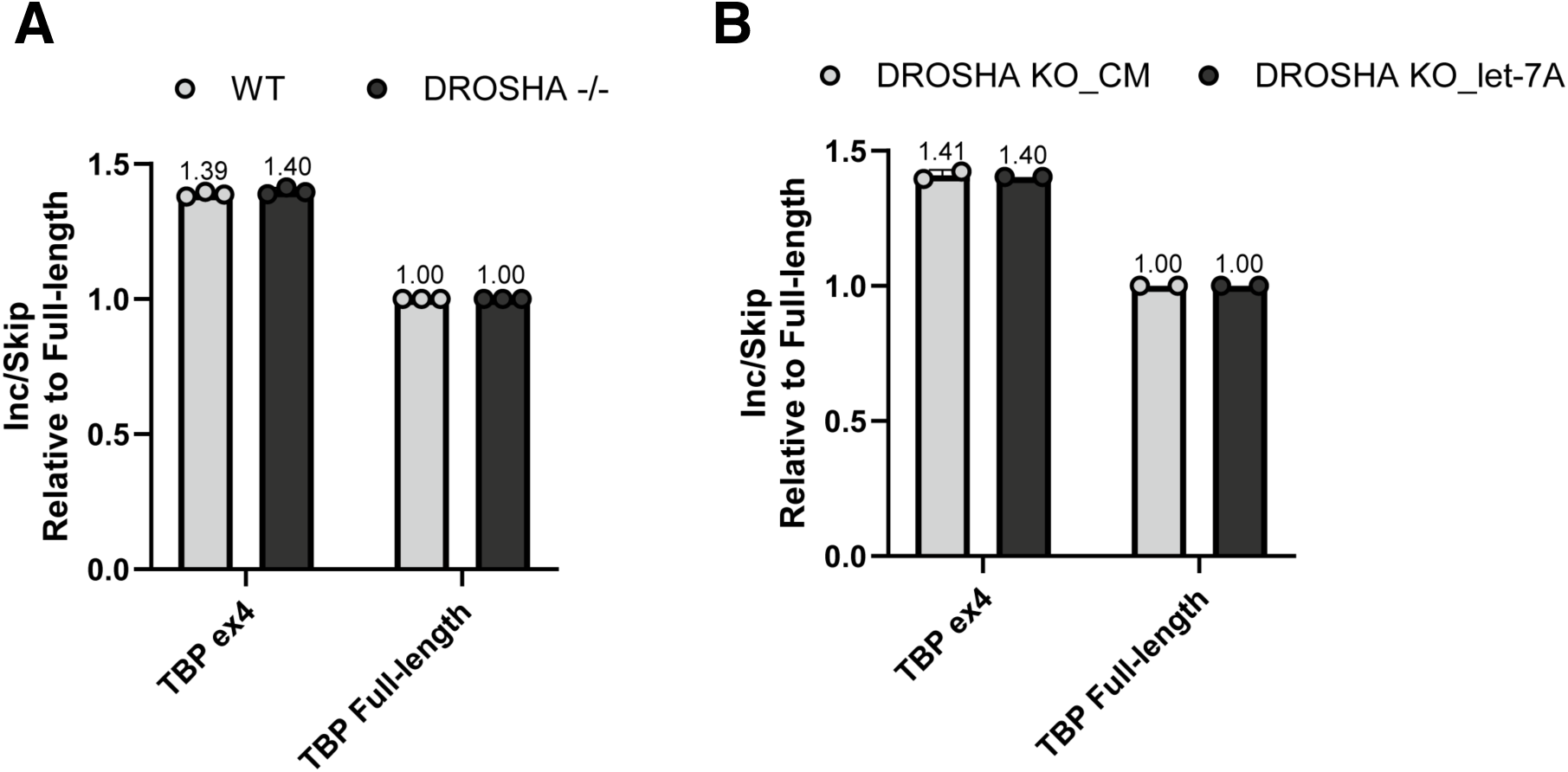
*Let-7* family miRNAs do not significantly affect splicing of a reference control gene, *TBP*. The RT-qPCR results from this assay, designed similarly to the *HMGA2* splicing assay, are plotted as splicing inclusion:skipping ratios, as shown in (**A)** for untreated WT and DROSHA-/- and (**B)** for DROSHA-/- transfected with *let-7a* miRNA mimic or control. Each condition’s respective mean value, calculated from independent biological replicates, are displayed above each condition shown.

